# Aphid infestation induces plant-sex-specific changes in floral chemistry and pollinator behaviour in *Silene latifolia*

**DOI:** 10.1101/2025.07.22.666187

**Authors:** Kaya B. Zill, Thomas Stegemann, Elisabeth Kaltenegger, Wolfgang Bilger, Tobias J. Demetrowitsch, Henry Berndt, Alexandra Erfmeier, Sybille B. Unsicker, Karin Schrieber

## Abstract

Pollinators share the complex information and resource landscape of their host plants with herbivores. Yet, how sap feeders affect floral attractiveness to pollinators remains poorly understood, despite the critical role of this tripartite interaction in natural and agricultural ecosystems. In dioecious plant species, which display pronounced sexual dimorphism, these intricate interactions may vary in magnitude and direction between females and males, with significant implications for plant population dynamics and species co-evolution. In this study, we examined how infestation by the oligophagous aphid *Brachycaudus lychnidis* affects sex-specific interactions among the dioecious plant *Silene latifolia* and its specialist moth pollinator *Hadena bicruris*. We exposed male and female plants to aphid herbivory and evaluated its effects on floral traits (visual cues, floral scent, and nectar chemistry) and pollinator behaviour. While aphid infestation affected some floral traits equally in both sexes and others more strongly in males or in females, we observed stronger declines in female attractiveness to pollinators, which were mainly linked to nectar compounds potentially acting as feeding cues or behavioural modulators. We discuss our results in the light of sexual selection and plant defence theory while emphasizing the complementarity of female and male traits in stabilizing this specialized plant-pollinator-herbivore system.

**Highlight:** Aphid infestation alters multiple visual and chemical floral traits in a plant sex-specific manner, leading to reduced attractiveness to moth pollinators in female plants, but not in males.

**Graphical Abstract:** Plant-sex specific effect of aphid infestation on floral traits (number, size, colour, scent composition, nectar quantity and composition) and pollinator behaviour.

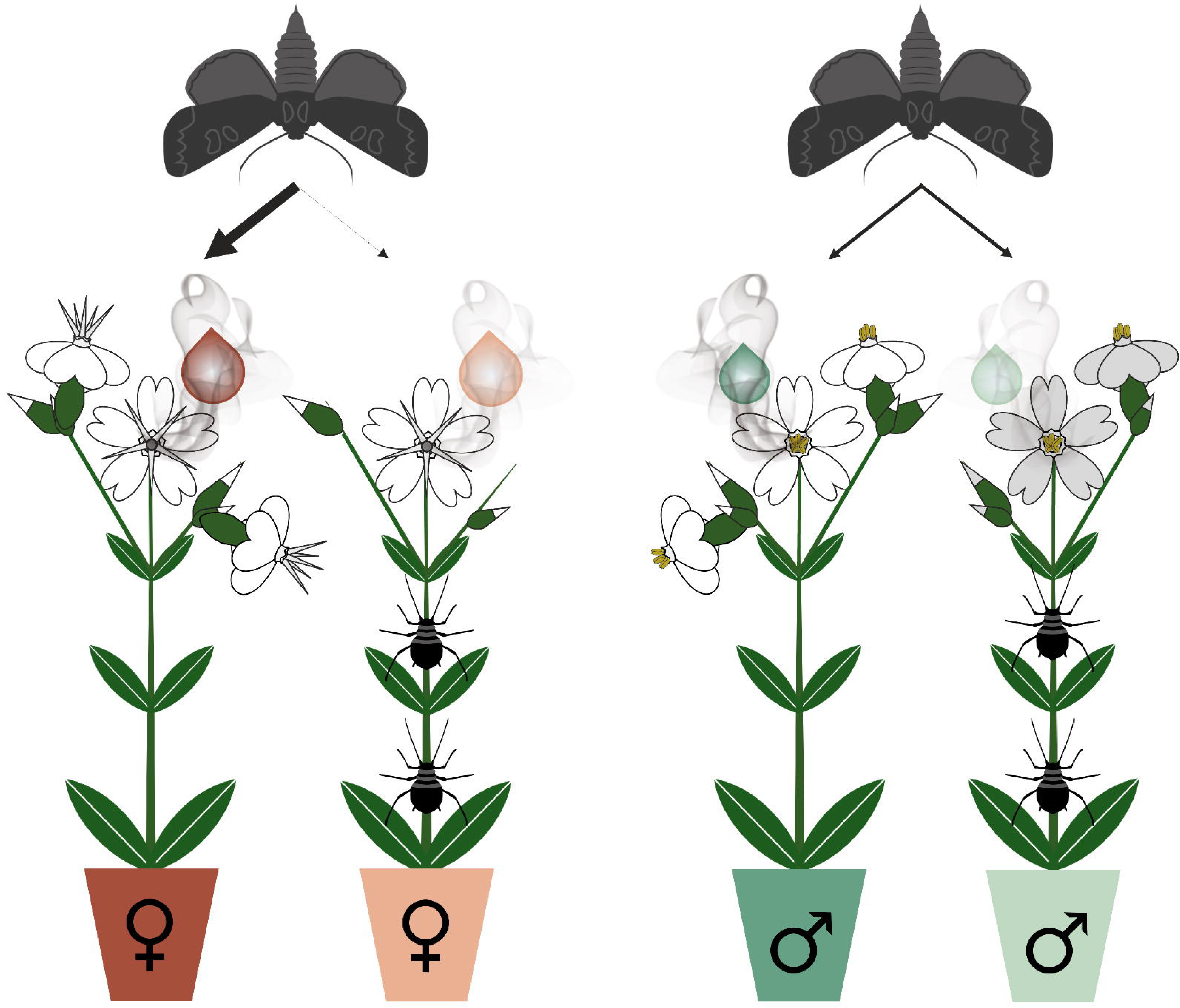

## 1. Introduction

Understanding plant × pollinator × herbivore interactions and their component species’ co-evolutionary dynamics has recently become a central goal in chemical ecology (Moreira *et al*., 2019; Kessler and Chautá, 2020; De-la-Cruz *et al*., 2022; Muola *et al*., 2022). The complex plant traits mediating interactions with pollinators and herbivores show considerable overlap, and we are beginning to understand the ecology of this shared trait space in systems involving chewing-biting herbivores. However, the impact of sap feeders on communication and resource exchange with pollinators remains largely unexplored, despite their critical role in the function of both natural and agricultural ecosystems (Morales-Hojas, 2017; Noriega *et al*., 2018).

Plants evolve diverse traits to attract pollinators and modulate their behaviour thereby maximizing their own fitness with minimal resource investment (Santamaría and Rodríguez-Gironés, 2015). Next to flowering phenology and visual signals (flower number, spatial arrangement, size, shape, hue, and colour luminance), floral chemistry plays a central role in ensuring these complex functions (Borghi *et al*., 2017; Trunschke *et al*., 2021; Wong *et al*., 2023). The emission of floral volatile organic compounds (VOCs) can trigger oriented movement of the receiver towards the emitter, and elicit landing, feeding as well as reproductive behaviour (Muhlemann *et al*., 2014). Such scent signals can be honest and guide visitors to a high-quality reward, or deceive them about the presence of food or mates, thereby exploiting their pollination service (Ito *et al*., 2021; Peakall, 2023). Nectar - the primary nutritional reward provided to pollinators - is a complex secretion composed of sugars, amino acids, lipids, proteins and diverse secondary metabolites (Nicolson, 2022), and thus not only a ready-to-use energy source for feeding animals. Plants may use nectar as a toolkit to manipulate the behaviour of their floral visitors. Secondary nectar metabolites can cause phagostimulation and satiety; improve learning capacity and memory; promote muscle performance, motivation and arousal; but also initiate sluggish and imprecise movements (Nepi *et al*., 2018; Mustard, 2020; Barberis *et al*., 2023). These various functions facilitate the exploitation of highly efficient pollination services while reducing energy investment in nectar sugars and amino acids. Although we are becoming increasingly aware of this beautiful complexity, there are still few studies that take a holistic view of the pollination syndrome, considering the combined roles of visual traits, floral VOCs, and nectar metabolites.

Pollinators universally share the complex information and resource landscape of their host plants with herbivores. There is now evidence for interactions among traits and genes controlling plant reproduction and defence strategies, leading to complex reciprocal co-evolutionary feedbacks (Kessler and Chautá, 2020; De-la-Cruz *et al*., 2022). Herbivores may directly reduce plant attractiveness to pollinators through aggression or the consumption of floral tissues and vegetative structures necessary for flower production (McCall and Irwin, 2006). Multiple floral traits have thus evolved to protect floral tissues from herbivory (Paul *et al*., 2022). Constitutive and induced plant defences may not only alter a plant’s suitability to herbivores, but also deter pollinators (Adler *et al*., 2012) or reduce resource allocation to pollinator attraction (Stamp, 2003). Specifically chemical plant traits are considered to mediate multiple other antagonistic and synergistic pleiotropic effects on pollinator attraction and herbivore defence (Kessler and Chautá, 2020). Defence compounds induced by herbivores can for example be allocated to nectar or pollen and deter or harm pollinators (Rivest and Forrest, 2020), while specific floral VOC attracting pollinators can be used as a feeding cue for florivores (Sasidharan *et al*., 2023). So far, insight into such complex tripartite interactions comes almost exclusively from chewing-biting herbivores. Aphids, which are among the most damaging plant pests (Dedryver *et al*., 2010), however, have been largely overlooked (but see Pareja et al., 2012). These exceptionally fast reproducing insects cause considerable stress in plants. They passively feed on phloem sap, thereby rapidly depleting their hosts of essential nutrients, altering their source-sink relationships, and excreting phytotoxins as well as phytopathogenic viruses with their saliva (Nalam *et al*., 2019; Twayana *et al*., 2022). Plants in turn, actively defend against aphids by changing the composition of sugars and amino acids in the phloem sap or loading it with deterrents, toxins or anti-digestive proteins (Nalam *et al*., 2019; Twayana *et al*., 2022). Floral traits attracting pollinators are specifically prone to the direct and indirect effects of aphid infestation, as aphids often feed close to reproductive structures on flowering stems. This proximity suggests a high potential for interference with pollinator-mediated reproductive processes, warranting closer empirical attention.

The eco-evolutionary link between pollinator attraction and herbivore defence becomes particularly apparent in dioecious species (Johnson *et al*., 2015). Plants that allocate male and female reproductive function to different individuals often exhibit complementary sexual divergence in both traits based on dramatic differences in resource demands for successful reproduction (Barrett and Hough, 2013; Johnson *et al*., 2015). Male success is predicted to largely depend on efficient pollen transfer and thus pollinator attraction, while female success relies on pollination, as well as on the production and defence of seeds, and their dispersal (Moore and Pannell, 2011). Consequently, females are predicted to allocate fewer resources to floral attractiveness but exhibit more efficient herbivory defence and tolerance as compared to males (Avila-Sakar and Romanow, 2012; Barrett and Hough, 2013). As recent research has produced contradictory evidence for these theories (Cornelissen and Stiling, 2005; Tang-Martínez, 2012; Sargent and McKeough, 2022), a central question remains: What are the sex-specific ecological consequences of parallel interactions with pollinators and herbivores in dioecious plant species? Addressing this question is key to understanding how co-selection by pollinators and herbivores contributes to the evolution of divergent reproductive strategies in dioecious plants.

A system that is ideal to study these complex interplays consists of the dioecious plant *Silene latifolia* POIR. (Caryophyllaceae), its nocturnal pollinator *Hadena bicruris* HUFN. (Noctuidae), and the aphid *Brachycaudus lychnidis* L. (Aphididae). The species involved in this system engage in highly specialized relationships. Researchers have only begun to uncover the sophisticated functional traits that have co-evolved to stabilize the cost-benefit ratio of their individual bipartite interactions (Blair and Wolfe, 2004; Dötterl *et al*., 2006; Labouche and Bernasconi, 2010; Schrieber *et al*., 2021). Insight into the ecology of the tripartite interaction, however, is lacking. Here, we exposed female and male *S. latifolia* plants to control and *B. lychnidis* infestation treatments to assess flowering phenology, spatial flower arrangement, flower colour, the chemical composition of floral scent and nectar as well as nectar quantity, and attractiveness to *H. bicruris*. We specifically aimed at characterizing i) sexual dimorphisms in floral attractiveness; ii) the effect of aphid infestation on floral attractiveness to pollinators; and iii) sex-specific differences in the susceptibility to aphid infestation and the quality and magnitude of responses to aphid infestation.

## 2. Methods

### 2.1 Study organism*s*

*Silene latifolia* (white campion, *Figure 1A*) is a short-lived dioecious plant species within the Caryophyllaceae family, which is native to ruderal habitats across Eurasia and North Africa. The plant exhibits a distinct moth pollination syndrome. Its white, large, and funnel-shaped flowers open at dusk to release a scent bouquet composed of more than 60 VOCs (Dötterl and Jürgens, 2005; Dötterl *et al*., 2005) to advertise large amounts of nectar rewards (Tew *et al*., 2021) with a mostly unknown chemical composition. Male and female *S. latifolia* are known to exhibit pronounced sexual dimorphisms in a broad set of floral traits. Observed differences in flower number, size, colour, total floral scent emission, and nectar volume as well as sugar content (Gehring *et al*., 2004; Waelti *et al*., 2009*a*; Delph *et al*., 2010; Schrieber *et al*., 2021)are largely in line with theory of sexual selection, i.e., higher trait values in male plants (Bateman, 1948).

**Figure 1:**
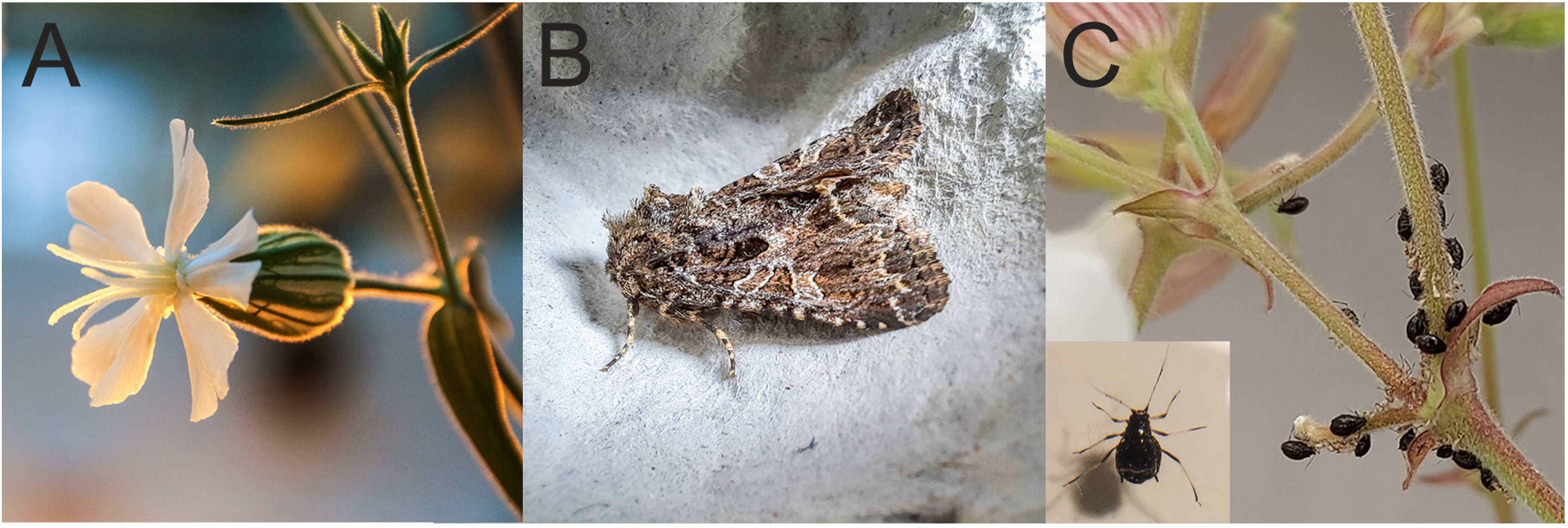
Plant-pollinator-herbivore study system. (A) *Silene latifolia* (female), (B) *Hadena bicruris* (male), (C) adult *Brachycaudus lychnidis* individual and colony feeding on flowering stems of S. latifolia.

*Silene latifolia* is readily visited by diurnal generalist insects and nocturnal moths, with *Hadena bicruris* (campion owl, *Figure 1B*) *-* a crepuscular moth within the Noctuidae family - being the most frequent and relevant pollinator species in Eurasia (Jürgens *et al*., 1996; van Putten *et al*., 2007; Schrieber *et al*., 2021). Together both species form one of only ∼16 worldwide known nursery pollination systems (Hossaert-McKey *et al*., 2010). Female moths pollinate female plants and oviposit on flower ovaries to provide their hatching larvae with nutritious seeds and shelter by developing fruits. The pollination services provided by male moths, together with the high egg and larval mortality caused by parasitoids and predators of *H. bicruris*, can compensate for the costs of seed consumption (Labouche and Bernasconi, 2010; Villacañas de Castro and Hoffmeister, 2020). Together these factors shift the outcome of the *Silene latifolia* × *Hadena bicruris* interaction for the plant towards the latter end of a seed predation-pollination continuum. A substantial fraction of VOCs in the floral scent of *S. latifolia* triggers antennal and behavioural responses in *H. bicruris* (Dötterl *et al*., 2006). While the function of floral scents in this intricate co-evolutionary relationship is well studied (Dötterl *et al*., 2006), the functional chemistry of nectar - the only reward and main food source provided to adult moths - remains unexplored beyond its basic sugar composition (Shykoff, 1997; Gehring *et al*., 2004; Witt *et al*., 2013).

A third species participating in this highly specialized system within Eurasia is the holocyclic aphid *Brachycaudus lychnidis (Figure 1C)*, which spends both the parthenogenetic and sexual part of its life cycle on *S. latifolia*. The aphid feeds mainly on the phloem sap of flowering stems.

### 2.2 Plant and insect cultivation

Seeds from five *S. latifolia* mother plants were collected in a population in Wiesengrund, Germany (51°41’17.1’N, 14°35’45.8’E) and experimentally outcrossed for two generations under greenhouse conditions (see Schrieber *et al*., 2019 for details). We cultivated 30 plants per maternal family individually in 360 mL pots filled with organic potting soil (Floragard TKS® 2 Instant Plus, DE) in a greenhouse at 25 °C / 15 °C and 60 % RH in a 16 h day / 8 h night cycle. When plants started flowering after 6 weeks, we selected two males and two females from each of the five maternal families (N = 20). The selected plants were transferred into 8 L pots filled with the above-mentioned substrate and pruned to a height of 15 cm to induce over-compensatory growth. Please note that *S. latifolia* is adapted to grazing/mowing and thus cutting is near-natural (Jessen *et al*., 2022). Moreover, cutting was necessary to induce the emergence of multiple shoots and sufficient flower production for the experiment. Plants were fertilized with 0.3 g Universol® Yellow (Everris-Headquarters, NL) dissolved in 200 mL of water every other week and prophylactically treated with sulphur vapor to prevent fungal infections as well as with biological pest control agents to prevent thrips and uncontrolled aphid infestation (agents: *Amblyseius cucumeris against thrips; Aphidius ervi, Aphidius colemani and Lysiphlebus testaceipes* against aphids Katz Biotech GmbH, DE). The preventive treatment with parasitoid wasps was stopped three weeks before controlled aphid infestation.

Adult *H. bicruris* moths were collected in different wild *S. latifolia* populations around Kiel, Germany (54°19′ N, 10°8′ O) in 2022 and kept together in ventilated, transparent 20 x 30 cm boxes equipped with egg cartons and tissue papers within a climate chamber (MLR-352H, PHC Corporation, Ōizumi-machi, JP) at 25 °C / 15 °C and 60 % RH in a 16 h day / 8 h night cycle. They were fed a honey-water solution (6 g organic flower honey, 14 mL distilled water) from plastic *S. latifolia* flowers. In addition, moths received one fresh male and female *S. latifolia* flower per day, as they did not reproduce without them. Individual flowers were harvested from a pool of food plants and presented in Eppendorf tubes filled with water. *Hadena bicruris* eggs, which were deposited all across the breeding boxes, were collected and placed on moist filter paper in individual ventilated plastic containers (50 mL). Upon hatching, larvae were provided with an artificial diet according to (Elzinga *et al*., 2002), which was additionally supplemented with 20 mL of *S. latifolia* seeds per 500 mL to increase acceptance. Pupated *H. bicruris* were moved to ventilated 50 mL containers filled with sterile sand and newly hatched moths were transferred into the above-mentioned breeding boxes and food supply for adults.

*Brachycaudus lychnidis* aphids were collected and identified according to (influentialpoints.com, 2024) in spring 2024 in different wild *S. latifolia* populations around Kiel, Germany (54°19′ N, 10°8′ O). They were cultivated in net cages on a balanced mixture of male and female *S. latifolia* plants from our study population in a climate chamber (HGC 1514 Module 7, Weiss Umwelttechnik GmbH, DE) at 23 °C and 55 % RH, in a 16 h day / 8 h night cycle. Food plants were watered and exchanged on demand.

### 2.3 Experimental setup and data acquisition

In mid-July 2024 at the age of 13 weeks, all selected plants were enclosed in 150 × 50 cm organza mesh bags and one individual per sex (male, female) × maternal family (I-V) combination was inoculated with 50 adult aphids *(Figure 2A, Supplementary Figure S1A-C)*. All aphid colonies showed an initial decline in population size of max. 10 % and switched to positive growth rates max. three days after the inoculation. Thus, our approach does not account for potential plant sex-specific host choice, early phase plant defences preventing colonization success, and indirect defence; but focuses on the effects of established aphid colonies having optimal growth conditions. Data on plant stress indicators, pollinator choice, visual flower traits, floral scent, and nectar *(Figure 2B-F)* were collected over a 40-day period following aphid inoculation *(Figure 2G)*.

**Figure 2:**
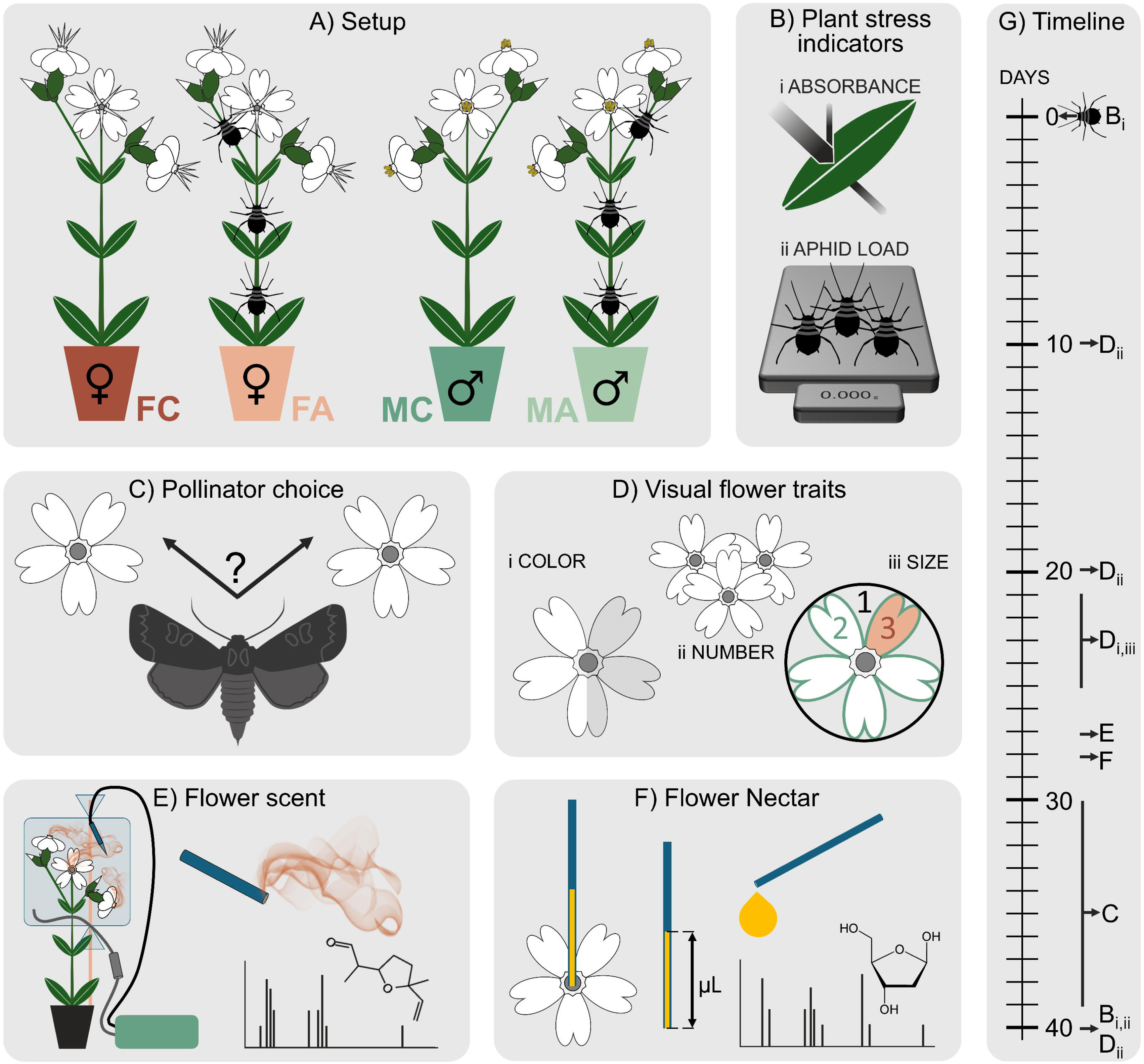
Overview of (A) the experimental setup (abbreviations: FC = control female, FA = aphid infested female, MC = control male, MA = aphid infested male); the data acquisition for (B) the plant stress indicators (i) absorbance by chlorophyll and epidermally located flavonoids and (ii) aphid load in terms of weight and number, (C) pollinator behaviour, (D) visual flower traits related to (i) color, (ii) quantity and (iii) display area, (E) floral scent, and (F) flower nectar; as well as (G) the time line for data acquisition after inoculation of plants with aphids.

#### 2.3.1 Plant stress indicators

To quantify the disruption of physiological homeostasis under aphid infestation, we optically assessed the chlorophyll concentration, and the absorbance by epidermally located UV-absorbing compounds, mainly flavonoids using the LEAF-STATE-ANALYZER LSA-2050 (Heinz Walz GmbH, DE) *(Figure 2B&G)* for all plants. As reference, data provided with the instrument for the lower leaf side of *Sedum telephium* was used. Data were acquired directly before and 40 days after aphid inoculation between 11:00 am and 12:00 noon in five leaves per plant individual. The selected leaves were evenly distributed along the vertical axis of the plants and well exposed to light (i.e., not significantly shaded by other leaves) and the measured area, located in the upper middle part of the lamina, was free of honeydew and aphids, which typically feed on stems. Immediately after these measurements, we counted the number of aphid individuals per size category (small: < 1.5 mm, medium: < 2 mm, large: > 2 mm) directly on the plant. Subsequently, the aphids were carefully washed off the plants with warm tap water, collected on filter paper and dried for 24 h at 25 °C, then for 24 h at 65 °C, and finally for 24 h at 105 °C to determine total aphid dry biomass per plant.

#### 2.3.2 Pollinator behaviour

Behavioural assays with *H. bicruris* were conducted under laboratory conditions 30-39 days after aphid inoculation *(Figure 2C&G)* between 8:00 pm and 0:30 am - the peak activity time of our captive moths. For the choice assays, freshly hatched adult moths were kept individually in ventilated, transparent boxes (15 × 12 × 11 cm) equipped with shelters and a feeding station featuring a honey-water solution as described in *chapter 2.2*. For each assay, the artificial diet was removed in the morning and replaced in the evening with two fully developed (male: fully dehisced anthers, female: fully expanded and papillous stigma; flower age 2-3 days), freshly picked flowers from the experimental plants, which were carefully fixed in lidless Eppendorf tubes filled with tap water. The two flowers were positioned on opposite sides of the boxes for maximum spatial separation *(Supplementary Figure S1D)* to provide the moths with the binary choice among the following four treatment groups: control females *versus* aphid infested females, control males *versus* aphid infested males, control females *versus* control males, aphid infested females *versus* aphid infested males. The moth individuals used in these behavioural tests (N = 11) differed in age and sex (5 females, 6 males; moth sex was accounted for in the statistical analysis) and were tested one to two times in each of the four treatment groups (depending on flower availability) using a fully randomized design that resulted in a total of 61 trials. Each night, up to eight moths were used, with two observers monitoring four moths each. The boxes were evenly lit with dim red LED lamps to allow observation without confounding the trials (the visual system of moths cannot capture red light). Three behavioural parameters were quantified for each of the two flowers: (i) the number of visits, with a visit starting and ending with physical contact to the flower; (ii) the number of feeding attempts per flower, with an attempt starting/ending by unrolling/rolling up and inserting/removing the proboscis into/from the corolla tube; and (iii) the total time spent with any physical contact on the flower. These three parameters were assessed for the first four visits within each trial. We deliberately decided against first-visit choice tests, as these give chance undue weight and moth individuals frequently revisit individual flowers or probe several flowers before feeding extensively on one under natural conditions (personal observation made in Schrieber *et al*., 2021). Female moths did not oviposit on the flowers offered in the choice test because they did not mate in isolation.

#### 2.3.3 Visual flower traits

We measured a broad set of visual flower traits at the plant individual and single flower level throughout the experiment *(Figure 2D&G)*.

Flower colour was quantified using a digital image transformation method that accounts for natural light conditions and the visual system of the pollinator based on (Troscianko and Stevens, 2015). Images were acquired 21-25 days after aphid inoculation for three to six flowers (depending on availability) from all plant individuals. They were taken outdoors in an open area, free from shadows cast by trees or buildings, during the last hour of daylight on rain-free days to match the natural light conditions perceived by *Hadena* (Bopp and Gottsberger, 2004). A fully developed but fresh flower (age 2-3 days) was plugged into a custom-made black ethylene vinyl acetate platform, which featured two reflectance standards (PTFB 10% and Spectralon 99%, Labsphere, UK) along with a size reference *(Supplementary Figure 1E)*. Images were captured using a modified Samsung NX1000 camera (Suwon, KOR) with its sensor filter removed, to achieve full-spectrum sensitivity (300 – 1000 nm). The camera was equipped with a UV-sensitive lens (Nikon EL 80 mm, JP). To capture both visible and ultraviolet (UV) light, we took one photo with a UV/IR blocking filter (UV/IR Cut, transmitting 400-700 nm, Baader Planetarium, DE) and another one with a UV pass plus IR blocking filter (U-filter, transmitting 300-400 nm, Baader Planetarium, DE) using a custom-made filter slide mount. All photos were taken with an aperture of 5.6, and ISO of 800, whereas the shutter speed was adjusted to the available light conditions to receive well-exposed photos. Post-processing was carried out using the Multispectral Image Calibration and Analysis (MICA) Toolbox plugin (Troscianko and Stevens, 2015) within ImageJ 1.47 t (Rueden *et al*., 2017). We selected the flower silhouette for colour analysis. The images were linearized to correct the camera’s non-linear light response and standardized based on the two reflectance standards to compensate for variations in natural lighting. We fitted the multispectral images to a mathematical model incorporating (i) the spectral sensitivity of our Samsung NX1000-Nikkor EL 80 mm 300-700 nm camera (data derived from Troscianko and Stevens, 2015); (ii) the spectral sensitivities of the three photoreceptors in the moth compound eye (data derived from Johnsen *et al.,* 2006); and (iii) the spectral composition of sun light during dusk (data derived from Johnsen *et al.,* 2006). For further details see Schrieber *et al*. (2021). The outcomes of these analyses are the percentage of green, blue and UV-A dusk sun radiation reflected by the flower, which are perceivable by a moth’s visual system.

Flower area was measured from the same images for three to six flowers per plant individual. We assessed the exact area covered by a single, well developed petal limb, the area covered by the exact flower silhouette and the area covered by the circular perimeter *(Figure 2D, 1 = flower perimeter, 2 = flower silhouette, 3 = petal silhouette)*. The circular perimeter reflects the flower’s maximum extension, but not its visual impact on pollinators due to natural variation in the shape of single petals, while the area covered by the exact flower silhouette may represent what pollinators see but not the plant’s investment into flowers (petal overlap), which can be captured by individual petal area.

The number of opened flowers and flower buds per plant individual were recorded at three time points during the experiment (10, 20 and 40 days after aphid inoculation, *Figure 2D&G*) and corrected for the flowers removed for other analyses. Directly after the removal of aphids (40 days after aphid inoculation), plant aboveground parts were cut off directly on top of the soil and separated into shoots (without leaves), and flowers. Plant parts were dried individually in paper bags for 24 h at 65 °C followed by an additional 24 h at 105 °C to determine dry biomass. During data acquisition for pollinator behaviour, flower colour, and nectar rewards, we collected the harvested flowers in plant individual-specific bags to assess their dry biomass as described above and added it to the total flower dry biomass.

#### 2.3.4 Flower scent

At day 27 after the aphid inoculation, floral VOC emission was collected for all experimental plants during the peak emission in *S. latifolia* between 10:00 pm (sunset) and 1:00 am (Dötterl *et al*., 2005) using a mobile push-pull system as described in Berkum *et al*., (2024) . Up to three fresh and fully developed flowers per plant (age 2-3 days) from different shoots were enclosed in 35 cm × 31 cm PET bags (RUBIN® Bratschlauch, DE), which were closed at both ends with cable ties *(Figure 2E)*. Compressed air entered the system after passing through an activated charcoal filter on the lower side of the bag at a flow rate of 1 L min^-1^ and was pulled out at the top through an adsorbent Poropak filter (Alltech, USA), using a vacuum at the rate of 0.6 L/min. After VOC collection, the filters were eluted with 200 μL dichloromethane (Sigma Aldrich, DE) containing 10 ng/μL of nonyl acetate (Sigma Aldrich, DE) as an internal standard. As the headspace of treated plants also included aphids, we additionally determined which compounds were exclusively released by *Brachycaudus*. We enclosed 60 aphids (20 small: < 1.5 mm, 20 medium: > 1.5 mm, 20 large: > 2 mm) from our laboratory population in the above-described PET bags to collect and analyse their volatile emissions as described for the plants with three replicates.

Samples were analysed on a gas chromatograph (TRACE GC1310, ThermoFisher Scientific, DE) equipped with a TraceGOLD TG-5MS column (30 m × 0.25 mm × 0.25 µm + 5 m; ThermoFisher Scientific, DE) and coupled with a mass spectrometer (TSQ Duo MS, ThermoFisher Scientific, DE). A sample volume of 1 µL was injected with a split ratio of 1:10 at 250 °C and migrated with a helium-flow of 1 mL min^-1^. The transfer line was set to 270 °C and the ion source of the mass spectrometer was operated at 230 °C and 70 eV for ionization. The temperature program started at 40 °C for 2 min, ramped to 180 °C at a rate of 6 °C min^-1^, ramped to 300 °C at a rate of 60 °C min^-1^, and finished with a 5 min hold time at 300 °C. Mass spectra of separated compounds were acquired in full scan mode between 33 and 350 m/z. An alkane standard mix (C_8_-C_20_, Sigma Aldrich, DE) was analysed under the same conditions to calculate linear retention indices (RI) for the targeted compounds. The data processing was performed with Chromeleon^TM^ 7.3 (ThermoFisher Scientific, DE). The putative identity of compounds was determined via comparisons of mass spectra and RI’s with databases (NIST Mass Spectral Search Program Version 3.0, Smith *et al*., 2004) and (Dötterl *et al*., 2005). The identities were assigned according to the following criteria: match factor to the NIST databank >750 and RI difference < 10. We selected all VOCs for the statistical analyses that could not be clearly identified as contaminations. The base peak in the mass spectrum of these VOC was used as quantitation ion. Additionally, two secondary ions (confirmation ions) were selected per compound to ensure correct peak assignment *(Supplementary Table S1A)*. The relative compound abundance was quantified by dividing the peak area (based on individual quantitation ion) of each volatile by the peak area of the internal standard nonyl acetate (quantitation ion 43, confirmation ions 56, 70). To account for variation in flower number in the headspace, relative scent abundance was normalized per flower. In addition to this semi-quantitative approach, we created another dataset to focus on qualitative differences only by dividing the peak area of each VOC by the total peak area of all compounds.

#### 2.3.5 Nectar quantity and chemistry

We measured the nectar volume for the flowers that have been harvested for the colour and size analyses, directly after they have been photographed. All nectar was extracted from the base of the ovary / the base of the stamens and anthopophor into 1 µL microcapillary tubes (Minicaps NA-HEP, Hirschmann Laborgeräte, DE). The length of the nectar column was measured with a calliper to determine the exact volume.

Nectar chemistry was addressed based on a standardized nectar volume via Ultra High Performance Liquid Chromatography - Quadrupole Time of Flight - Electro Spray Ionization - Tandem Mass Spectrometry (UHPLC-ESI-qTof-MS/MS). For these analyses, we additionally collected all nectar from three fresh, fully developed flowers per plant individual using 1 µL microcapillary tubes on day 28 after aphid infestation. After determining the exact nectar volume as described above, each microcapillary was rinsed with 30 µL of HPLC grade water (Agilent, USA) into a 0.5 mL Eppendorf tube, in which all nectar from one plant individual was pooled. The samples were then vortexed, snap-frozen in liquid N_2_ and stored at −80 °C. Samples underwent a modified simultaneous metabolite, protein, and lipid extraction (SIMPLEX) as described in (Schrieber *et al*., 2023) to obtain both, hydrophilic and lipophilic nectar metabolites. After defrosting and vortexing, 0.1 mL of nectar solution was mixed with 0.5 mL of ultra-pure methanol (Carl Roth, GE) and 4 mL of methyl-tert-butylether (LC-grade, Carl Roth, GE). Samples were incubated for 30 min at 25 rpm in an end-over-end shaker before phase separation was induced by adding 0.5 mL of ultra-pure water (Carl Roth, GE). After 10 min of incubation, samples were centrifuged at 4000 *g* and 4 °C for 10 min and the upper lipophilic phase was collected. The lower hydrophilic phase was used as a template to repeat the above-described protocol for the extraction of hydrophilic metabolites. Extracts were re-suspended in 0.5 mL of either ultra-pure water and ultra-pure methanol (hydrophilic extracts: 50 / 50, v / v, Carl Roth, GE) or ultra-pure isopropanol and LC-grade chloroform (lipophilic extracts: 3:1, v / v, Carl Roth, GE). The dilution of the resulting nectar stock solutions for analyses via UPLC-qTof-MS/MS was then adjusted to differences in the volume of extracted nectar among plant individuals. Hence, the volume of analysed nectar was standardized for these analyses.

For UHPLC-ESI-qTof-MS/MS analyses, 2 µL of extract were injected at 30 °C into a UHPLC (Nexara, Shimadzu, DE) containing a C_18_ Column (100 × 2.1 mm, 1.7 µm, Luna Omega Polar, Phenomenx, DE). The solvent system was composed of (A) water with 0.1% formic acid, and (B) acetonitrile with 0,1% formic acid (all LC-grade, Carl Roth, DE). The gradient program started at a flow rate of 0.3 mL min^-1^ with 5 % B, was then raised over 20 min to 95 % B, and finally kept at 95 % B for an additional 5 min. Afterwards, the column was re-equilibrated at 5% B for 6 min. Chromatographically separated analytes entered a qTOF-MS/MS (x500R, Sciex, DE) with the following parameters: ion source gas 45 psi (nitrogen), curtain gas 35 psi (nitrogen), CAD gas 7 psi (nitrogen), electrospray ion source temperature 500 °C, ion spray voltage 5500 V. The MS acquisitions parameters were: MS1 scans at 100-1000 m/z with a data-dependent MS2 trigger having a collision energy (CID) of 40 eV and a scan range of 50-1000 m/z. The acquired data were then converted to .mzML format and further processed in mzMine (Schmid *et al*., 2023). Metabolic features with a maximum m/z difference of 0.001 amu and maximum retention time difference of 0.1 min were bucketed and treated as identical features in the following analysis. For feature annotation, the acquired MS/MS spectra were compared against public mass spectral libraries from Global Natural Products Social Network (GNPS) (Wang *et al*., 2016) and the MassBank (MassBank consortium and its contributors, 2024) using the NIST search algorithm (cosine similarity of 0.6) and the MassBank formula (cosine similarity of 0.5). Features that remained unidentified were further annotated using COSMIC and CSI:FingerID according to the SIRIUS software package (Hoffmann *et al*., 2022).

#### 2.3.6 Statistical analyses

All statistical analyses were implemented in R 4.4.1 (R Development Core Team, 2024). Plant stress indicators, spatial flower traits, flower colour traits, and floral rewards were analysed as responses in (generalized) linear mixed effects models - (G)LMMs, which included the aphid infestation treatment, plant sex and the interaction among both variables as predictors. Highly correlated traits were combined into a single response and analysed within one model. These models included the different types of summarized response traits as a factorial co-predictor, for which all possible interactions with sex and aphid infestation treatment were tested according to the scheme *response at plant individual level (level of summary) ∼ cofactor:* leaf chlorophyll content (on day 0 and 40 of the experiment) ∼ time point; leaf epidermal light absorption by flavonoids (on day 0 and 40 of the experiment) ∼ time point; number aphids (belonging to the size category small, medium and large) ∼ aphid size category; flower number (on day 10, 20 and 40 of the experiment) ∼ time point; dry biomass (of a single flower, all flowers, and flowering stems) ∼ biomass component; display area (of a single petal limb, all petal limbs, and the full corolla expansion) ∼ flower component; flower light reflection (in the green, blue and UV-A range) ∼ colour quality. The remaining traits (nectar volume) were acquired across several flowers per plant individual and analysed in this high resolution (i.e., not based on means). Consequently, all models had repeated measurements on the plant individual level and thus included a random effect for the plant individual. All models were checked for violations of their underlying assumptions using tests and plots from the R-package ‘DHARMa’ (Hartig, 2022) and were adjusted with regard to error family, dispersion or zero-inflation, if necessary. An overview of all model parameters is provided in *Table 1*. Based on these models, we calculated type III ANOVA tables (based on Wald- χ² tests) while setting all contrasts to sum-zero coding using the R-package “car” (Fox and Weisberg, 2019). If factorial predictors were involved in significant interactions, we additionally calculated post-hoc contrasts among the estimated means of their levels within levels of other factors involved in the respective interaction (R-package “emmeans”; Lenth, 2024)).

**Table 1:** Overview of statistical analyses investigating the effects of aphid infestation treatment (atm) with *Brachycaudus lychnidis* and *Silene latifolia* plant sex on plant stress indicators, behaviour of the pollinator *Hadena bicruris* and plant traits involved in communication and resource exchange with pollinators. The table provides an overview of the model structures with regard to the included responses (column 1), predictors and random effects (row 1); as well as further model characteristics such as response transformations (log = natural logarithm, none = none), model types (GLMM, LMM, LM, PERMANOVA), the assumed error families (B = binomial, BB = betabinomial, G = Gaussian, NB = overdispersed Poisson with quadratic parametrization, P = Poisson), and additional zero-inflation / dispersion formulas (Z / D∼ = account for access of zeroes in Poisson data / account for predictor-specific variance inhomogeneity, none = none). Moreover, the table provides an overview of p-values from type III ANOVAs (based on Wald χ²-test for (G)LMMs, F-tests for LMs, and permutation statistics with pseudo-F estimation for PERMANOVA). Significant predictor terms are highlighted with bold lettering. Main effects and lower order interaction terms involved in significant (higher order) interactions are marked with “ii”. Effects that were not tested are marked with ”-“. Please note that (†) the identity of cofactors for each response is described in detail in chapter 3.3.6; and (††) that in pollinator behaviour models, the actual test whether pollinators prefer control (C) over aphid infested (A) plants or female (F) over male (M) plants is tested by assessing whether the model intercept differs from 0.5. This represents the deviation from an equal choice and is not expressed as an interaction term with other predictors. Group-specific patterns are thus evaluated through post-hoc comparisons of the estimated probabilities for the respective factor levels, which are described in the respective text and figures.. SQ = semi-quantitative approach, QO = qualitative only approach.

|  | (1)<br>Sex | (2)<br>Atm | (3)<br>Cofactor† | (4)<br>Sex ×<br>Atm | (5)<br>Sex ×<br>Cofactor† | (6)<br>Atm ×<br>Cofactor† | (7)<br>Sex ×<br>Amt. ×<br>Cofactor† | (8)<br>Sex<br>moth | Random<br>effects |
| --- | --- | --- | --- | --- | --- | --- | --- | --- | --- |
| **Aphid number**<br>None, GLMM, NB, none | 0.328 | - | **0.000** | - | - | 0.207 | - | - | Plant<br>individual |
| **Aphid weight**<br>None, LM, G, none | 0.535 | - | - | - | - | - | - | - | - |
| **Leaf biomass**<br>None, LM, G, none | **0.035** | 0.516 | - | 0.397 | - | - | - | - | - |
| **Chlorophyll**<br>Log, LMM, G, D~(1-7) | **0.000** | 0.774 | **0.000** | 0.206 | 0.530 | 0.495 | 0.938 | - | Plant<br>individual |
| **Absorption by**<br>**Flavonoids**<br>None, LMM, G, none | **0.028** | 0.637 | **0.000** | 0.167 | 0.421- | 0.539 | 0.448 | - | Plant<br>individual |
| **Pollinator choice C-A**<br>None, GLMM, B, none | **0.000** | **0.072** | 0.986 | †† | 0.301 | †† | †† | **0.097** | Moth<br>individual,<br>trial |
| **Pollinator choice F-M**<br>None, GLMM, BB, none | 0.380 | 0.844 | 0.937 | †† | †† | 0.971 | †† | 0.572 | Moth<br>individual,<br>trial |
| **Flower number**<br>None, GLMM, P, Z~ I<br>D~(1) | **0.000<sup>ii</sup>** | **0.082<sup>ii</sup>** | **0.000<sup>ii</sup>** | 0.238<sup>ii</sup> | 0.757<sup>ii</sup> | **0.000<sup>i</sup>** | **0.012** | - | Plant<br>individual |
| **Flower biomass**<br>None, LMM, G, D~(3) | **0.000<sup>ii</sup>** | **0.001<sup>ii</sup>** | **0.000<sup>ii</sup>** | 0.123<sup>ii</sup> | **0.000<sup>ii</sup>** | **0.000<sup>i</sup>** | **0.000** | - | Plant<br>individual |
| **Flower colour**<br>None, LMM, G, D~(1,3,5) | 0.394<sup>ii</sup> | **0.000<sup>ii</sup>** | **0.000<sup>ii</sup>** | **0.027** | 0.977 | **0.016** | 0.6908 | - | Plant<br>individual |
| **Flower area**<br>None, GLMM, NB, D~(1-7) | 0.348 | 0.092 | **0.000** | 0.645 | 0.124 | 0.942 | 0.53 | - | Plant<br>individual |
| **Scent metabolome SQ**<br>PERMANOVA (Bray-Curtis) | 0.568 | **0.087** | - | 0.472 | - | - | - | - | - |
| **Scent metabolome QO**<br>PERMANOVA (Bray-Curtis) | 0.625 | 0.231 | - | 0.493 | - | - | - | - | - |
| **Nectar volume**<br>None, LMM, G, none | **0.000** | 0.554 | - | 0.225 | - | - | - | - | Plant<br>individual |
| **Nectar metabolome (LC)**<br>PERMANOVA (Bray-Curtis) | **0.000** | 0.204 | - | 0.308 | - | - | - | - | - |

The choice tests were analysed with two binomial models testing (i) preferences for a specific aphid infestation treatment within sexes (assays: control female *versus* aphid-infested female, control male *versus* aphid-infested male); and (ii) preferences for a specific plant sex within aphid infestation treatments (assays: control female *versus* control male, aphid infested female *versus* aphid-infested male). Model i / ii included the behavioural parameter in control relative to aphid infested flowers, i.e., the proportion of visits, feeding attempts or time spent in control relative to infested flowers (i) or female relative to male flowers, i.e., the proportion of visits, feeding attempts or time spent in control female to male flowers (ii) as binomial response and the flowers sexes (i) or the flowers aphid infestation treatments (ii) as predictors. Further predictors included in both models were the behavioural parameter type (visits, feeding attempts, time; all acquired in the same moth and trial), and its interaction with the previously mentioned model-specific predictor. In addition, both models included the sex of the moth as a cofactor for which only additive effects were tested, as sample sizes were not large enough to reliably test for all possible interactions. Moth individual and trial were set as random effects *(Table 1)*. Model checking, adjustment and downstream analyses with type III ANOVA were implemented as described above. Post-hoc contrast (R-package “emmeans”; Lenth, 2024) were used to test for a significant deviation of the binary outcome from the intercept of 0.5 (i.e., preference or avoidance) within the different plant sex × behavioural type combinations (i) or aphid infestation treatment × behavioural type combinations (ii).

The statistical approaches used for the analyses of the metabolome are identical for floral nectar and scent. In a first step, we performed a permutational analyses of variance (PERMANOVA) using the Bray-Curtis dissimilarities of the full metabolome datasets (n = 33 features in scent [for both, semi-quantitative and qualitative only data sets], n = 4,385 features in nectar assessed via UPLC-qTof-MS/MS) as the response distance matrix. We included aphid infestation treatment, sex, and their interactions as predictors and ran the models with 100,000 permutations (R-package “vegan” (Oksanen *et al*., 2024)). As previously implemented tests for multivariate homogeneity of group dispersions in the distance matrices among all treatment levels yielded no deviation for all datasets, the predictor effects assessed via PERMANOVA are based on differences in group centroids rather than differences in group dispersions (Anderson and Walsh, 2013). The multivariate group separations among plant infestation treatments and sexes were illustrated with Non-metric Multidimensional Scaling (NMDS) plots (R-package “vegan” (Oksanen *et al*., 2024)). To identify the compounds responsible for this separation, we used an approach that is comparable to a volcano plot (Cui and Churchill, 2003) and simultaneously allows the statistical assessment of interaction effects among plant sex and aphid infestation treatment. Instead of applying two-sided significance tests (t or u-tests), we entered all metabolites as responses into a linear model loop to test for the main and interactive effects of plant sex and aphid infestation treatment. Each metabolite ran with three alternative models as untransformed, squareroot-transfromed and log-transformed response. To check for violations of model assumptions and select the best of the three alternative models for each metabolite accordingly, we extracted the results from the Kolmogorov-Smirnov test for uniformity (normality of residuals) and Levene’s test for homogeneity of variance, which are provided for targeted model optimization in the R-package ‘DHARMa’ (Hartig, 2022). If model assumptions were violated in all three alternative models, the respective metabolite was not further analysed. The best fit models for the remaining metabolites were further analysed with type III ANOVAs based on F-tests (R-package “car” (Fox and Weisberg, 2019)) and the calculation of post-hoc contrasts (R-package “emmeans” (Lenth, 2024)) as described in the previous section. *P*-values were corrected for inflation of alpha errors by multiple testing (false discovery rate: FDR) according to (Benjamini and Hochberg, 1995). If a metabolites p-value in a given group comparison (F_C&A_/M_C&A_, T_F&M_/C_F&M_, FC/FT, MC/MT, FC/MC, FT/MT) was significant and the corresponding log_2_-fold change in the intensity of this metabolite was either ≤ −1 (halving of intensity in the first relative to the second listed group) or ≥ 1 (doubling of intensity in the first relative to the second listed group) it was considered to be modulated.

## 3. Results

### · 4.1 Plant stress indicators

After 40 days of inoculation with *B. lychnidis*, aphid load in terms of weight and number was equally high in female and male *S. latifolia* plants *(Table 1)*. Females had significantly higher trait values than males for leaf dry mass per plant individual (*females = 5.33 +/- 0.56, males = 3.50 +/- 0.56, p<0.05, F_2DF_= 5.28*), leaf chlorophyll content (*females = 27.5 +/- 1.37, males = 22.9 +/- 0.91, p<0.001,* χ*²_2DF_= 14.37)* and absorption by epidermal flavonoids (*females = 0.70 +/- 0.03, males = 0.62 +/- 0.03, p< 0.05,* χ*²_2DF_= 4.80),* but aphid infestation treatment had no effects on these traits *(Table 1)*. Moreover, leaf chlorophyll contents decreased *(p<0.001,* χ*²_2DF_= 60.34)*, whereas absorption by epidermal flavonoids increased *(p<0.001,* χ*²_2DF_= 49.47)* from the beginning towards the end of the experiment, regardless of sex and aphid infestation treatment *(Table 1*, column cofactor).

### 3.1 Effects of plant sex and aphid infestation on pollinator behaviour

A subset of our choice assays tested behavioural moth responses to single flowers from control versus aphid-infested plants, and whether these responses varied between male and female plants. The effects of aphid infestation treatment and plant sex on choice behaviour were marginally significant and highly significant, respectively *(Table 1; p= 0.072,* χ*²_2DF_= 3.23; p< 0.001,* χ*²_2DF_= 142.91)*. Post-hoc comparisons showed that only females, but not males, exhibited a highly significant preference for flowers from control plants over aphid-infested ones across all behavioural parameters *(Figure 3A; z-ratio female plants C-A: 3.36*** [visits], 3.23*** [feeding attempts], 4.19*** [time spent]; z-ratio male plants C-A: −0.08^ns^ [visits], - 0.24^ns^ [feeding attempts], −0.56^ns^ [time spent]).* In addition, we detected a marginally significant main effect of pollinator sex *(Table 1, Supplementary Figure S2, p=0.097,* χ*²_2DF_= 2.89),* indicating that the overall choice behaviour differed slightly between females and males. Post-hoc tests showed that only female but not male *H. bicruris* moths showed a significant preference for control over aphid-infested *S. latifolia* plants *(z-ratio female moth C-A: 2.45*, z-ratio male moth C-A: 0.16^ns^, Supplementary Figure S2)*.

**Figure 3:**
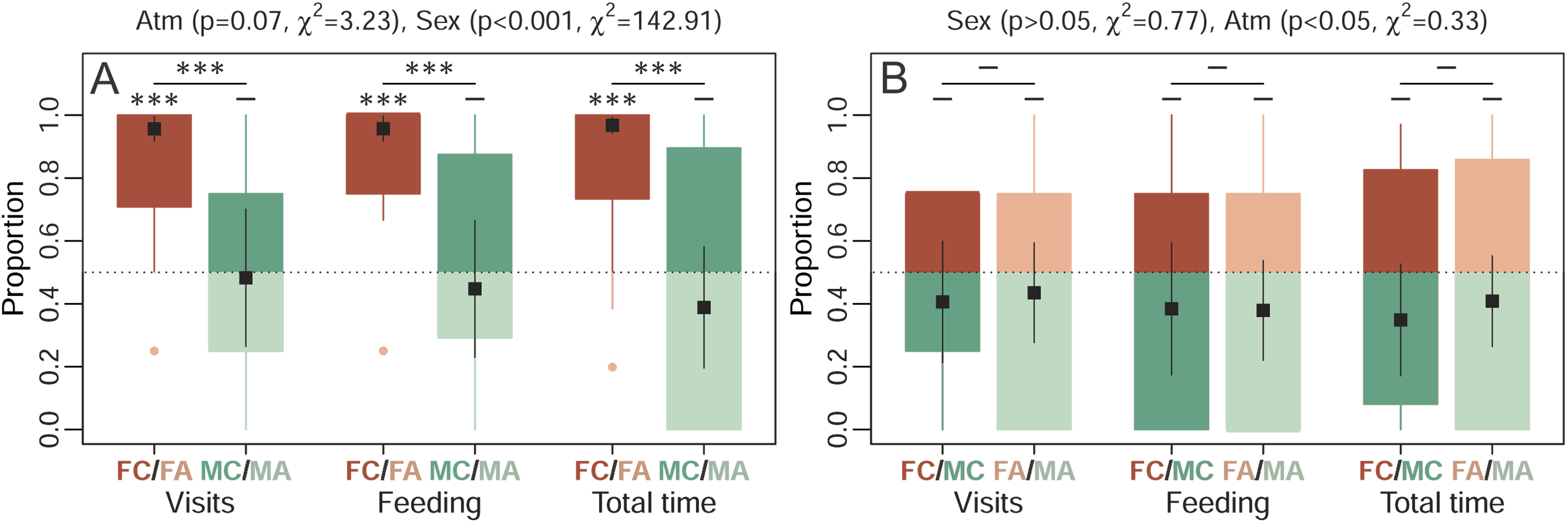
Effects of *Silene latifolia* plant sex and infestation by *Brachycaudus lychnidis* on the choice behaviour of the moth pollinator *Hadena bicruris* are analysed and illustrated as proportions of flower visits, feeding attempts and the total time spent between two offered flower types. (A) Illustrates preference (i.e., deviation from a binomial outcome of 0.5 marked by the dashed line) for control (C, deep coloured, outcome > 0.5) versus aphid infested plants (A, light coloured, < outcome 0.5) in *S. latifolia* females (F, red) and males (M, turquoise) (FC/FA: n=11, MC/MA: n=22). (B) Illustrates preference for female (F, red, outcome > 0.5) versus male (M, turquoise, outcome < 0.5) control (C, dark coloured) and aphid infested (A, light coloured) plants (FC/MC: n=19, FA/MA: n=9). Graphs combine Box-Whisker plots of raw data (box: IQR Q1-Q3, whiskers: smallest/largest values within Q1/Q3 - /+ 1.5 × IQR, points: outliers) with the marginal means +/- standard error (black squares with black lines) estimated for all groups by the (G)LMMs. Effects assessed via type III ANOVAs based on Wald χ² tests are denoted at the top of the plot, while the results of relevant post-hoc comparisons are denoted within the plots (significance levels: ***p<0.001, **p<0.01, *p<0.05, -: not significant). Asterisks printed on top of lines indicate differences of binomial outcomes among the connected treatment groups, while asterisks printed on top of individual boxes indicate significant deviations of the binomial outcome from 0.5 (i.e., preferences within a specific treatment group).

Another subset of our choice assays tested whether moths prefer single flowers from one plant sex over the other and whether this preference is modified by the aphid infestation treatment. The statistical analysis yielded no significant preferences of the moths for either plant sex and neither aphid infestation treatment nor their own sex shaped their choice *(Figure 3B, Table 1)*.

### 3.2 Effects of plant sex and aphid infestation on visual flower traits

Flower number was significantly shaped by the three-way interaction sex × infestation treatment × and infestation time *(Table 1, Figure 4A, p< 0.05,* χ*²_2DF_= 8.88)*. Post-hoc comparisons revealed that female plants suffered both earlier and stronger declines in flower number upon aphid infestation as compared to males *(female t-ratio C-A: 2.05^ns^ [10 days], 4.25*** [20 days], 9.26*** [40 days]; male t-ratio C-A: 0.40^ns^ [10 days], 1.36^ns^ [20 days], 4.74*** [40 days])*. Correspondingly, females produced significantly fewer flowers than males only in aphid infested plants and the magnitude of this effect increased over time *(control t-ratio F-M: -2.14^ns^ [10 days], −1.41^ns^ [20 days], −1.26^ns^ [40 days]; infestation t-ratio F-M: −3.84** [10 days], −4.31*** [20 days], −5.31*** [40 days])*. Reproductive biomass depended on the three-way interaction sex × infestation treatment × component *(Table 1, Figure 4B, p< 0.001,* χ*²_2DF_= 16.86)*. Post-hoc comparisons revealed that the component single flower mass was not affected by the infestation treatment *(t-ratio C-A: 1.17^ns^ [female], 1.14^ns^ [male])* but generally higher in females than males *(t-ratio F-M: 6.22*** [control], 5.76*** [infested])*. In contrast, the component total flower mass declined severely under aphid infestation, specifically in females *(t-ratio C-A: 12.89*** [female], 7.26*** [male])* and was accordingly higher in females only in controls but not under aphid infestation *(t-ratio F-M: 5.22*** [control], −0.41^ns^ [infested])*. The mass component of flowering stems was not affected by aphid infestation *(t-ratio C-A: 2.73 ^n.s^ [female], 0.86 ^n.s^ [male*]), and plant sex *(t-ratio F-M: 2.56^ns^ [control], 0.68^ns^ [infested])*. Flower light reflectance *(Table 1, Figure 4C)* was significantly shaped by the two two-way interactions sex × aphid infestation treatment (*p< 0.05,* χ*²_1DF_= 4.91)* and aphid infestation treatment × colour type (*p< 0.05,* χ*²_2DF_= 8.27*). Post-hoc comparisons showed that the aphid-infestation treatment reduced floral light reflectance in the green and blue spectrum only in male, but not in female plants *(female t-ratio C-A: 1.39^ns^ [green], 1.69 ^ns.^ [blue], 0.11^ns^ [UV]; male t-ratio C-A: 3.00* [green], 3.22* [blue], 2.09^ns^ [UV]).* Flower area was neither affected by plant sex, aphid infestation treatment or their interaction *(Table 1)*.

**Figure 4:**
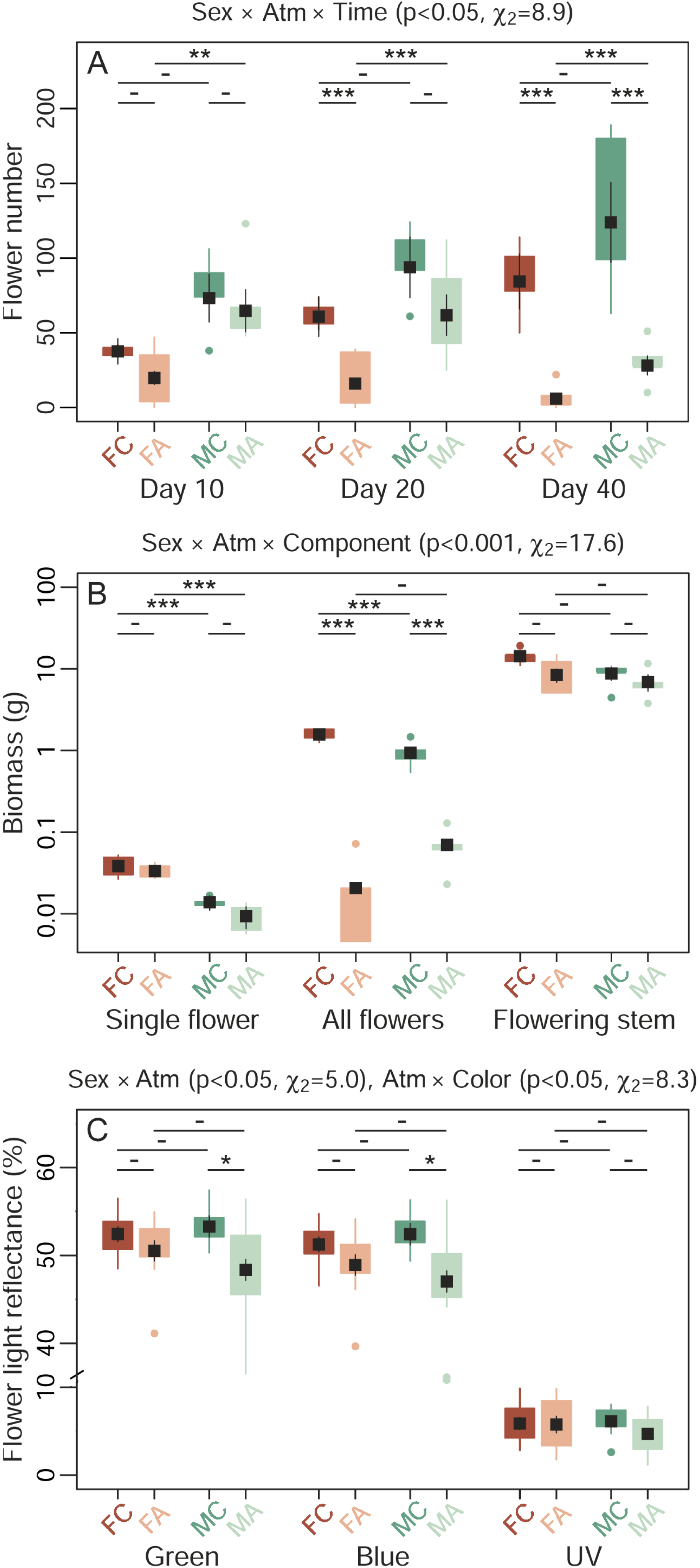
Combined effects of *Silene latifolia* plant sex and *Brachycaudus lychnidis* aphid infestation treatment on (A) flower numbers, (B) reproductive biomasses, and (C) flower colours. The graphs combine Box-Whisker plots of raw data (box: IQR Q1-Q3, whiskers: smallest/largest values within Q1/Q3 -/+ 1.5 × IQR, points: outliers) for female control (FC, dark red, n = 5_(A,B)_/14_(C)_), female infested (FA, light red, n = 5_(A,B)_/7_(C)_), male control (MC, dark turquoise, n = 5_(A,B)_/13_(C)_), and male infested (MA, light turquoise, n = 5_(A,B)_/13_(C)_) plants with the marginal means +/- standard error (black squares) for these groups estimated by the (G)LMMs. Significant effects detected via type III ANOVAs based on Wald χ² tests are denoted at the top of the plot, while the results of relevant post-hoc comparisons are denoted within the plots (significance levels: ***p<0.001, **p<0.01, *p<0.05, - not significant).

### 3.1 Effects of plant sex and aphid infestation on the composition of floral scent

We detected a total of 33 VOCs in the floral headspace of *S. latifolia*, with values normalized per flower. Of these 32 could be putatively identified via matches of RIs and mass spectra with databases *(Supplementary Table S1A)*. A large fraction of these VOCs have been previously described in *S. latifolia* (Dötterl *et al*., 2005; Mamadalieva *et al*., 2014) and of these, 16 have been shown to elicit antennal and/or behavioural responses in *H. bicruris* (Dötterl *et al*., 2006).

The PERMANOVA on the semi-quantitative VOC dataset detected a marginally significant main effect of aphid infestation treatment on the composition of these floral VOCs *(Table 1, p < 0.087, F_1DF_ = 2.30, R² = 0.13)* but no divergence among sexes or interactive effects. This is in accordance with both, the group dispersion visualized in the NMDS ordination plot *(Figure 5A)* and the outcomes of the model loop *(Figure 5B, Supplementary Table S1A)*. Without FDR correction, the loop identified eight compounds shaped by the main effect of treatment with reduced abundance in aphid infested as compared to control plants. These included five compounds known to elicit behavioural responses in *H. bicruris* according to Dötterl *et al*. (2006) -benzaldehyde, trans-β-ocimene, lilac aldehyde A, lilac aldehyde B+C, and lilac alcohol D- as well as two of their degradation products, lilac alcohol formate A and lilac alcohol formate B. One compound which was not tested for behavioural effects on *H. bicruris* so far - phenethyl benzoate - was significantly modulated by the interaction plant sex × aphid infestation treatment and decreased under aphid infestation in female plants but not in males. When accounting for inflation of alpha errors, all these effects were non-significant. These effects were detected despite the limited sample size, and might become more pronounced with additional replication. Multivariate analyses of qualitative differences only, yielded no significant effects of aphid infestation treatment and plant sex *(Table 1)*.

**Figure 5:**
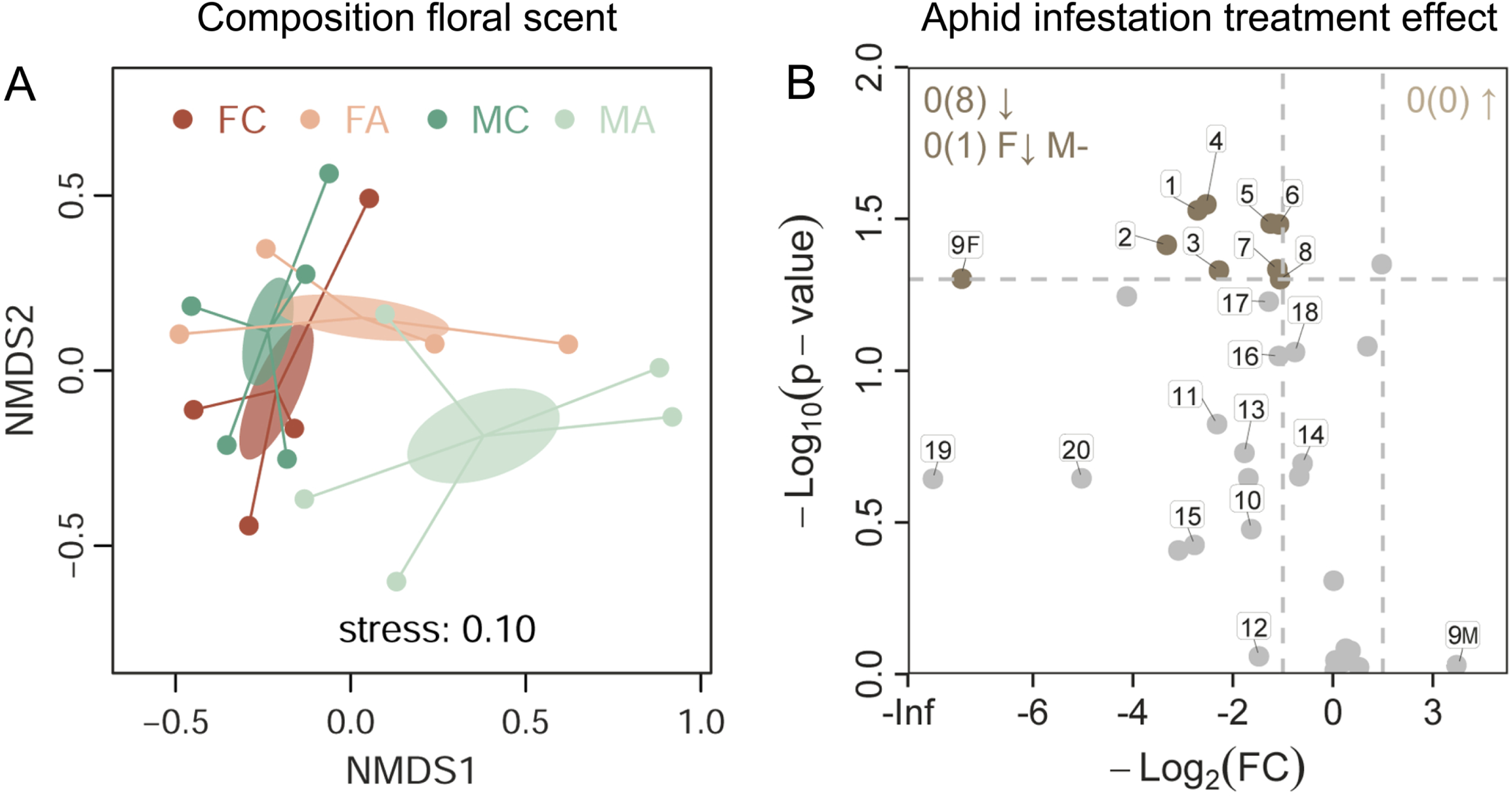
Combined effects of *Brachycaudus lychnidis* aphid infestation treatment and plant sex on floral headspace VOC composition in *Silene latifolia.* (A) The NMDS ordination for female control (FC, dark red, n = 4), female infested (FA, light red, n = 4), male control (MC, dark turquoise, n = 5), and male infested (MA, light turquoise, n = 5) plants shows plant individual scores (points), distances to group centroids (spiders), and their standard error (ellipses). (B) Volcano plot summarizing the results of model loops, i.e., the main and interactive effects of plant sex and aphid infestation treatment on the abundance of individual metabolites. Metabolic features modulated by aphid infestation are highlighted (log_2_ fold change <-1 or >1 and uncorrected [after FDR correction all effects non-significant] p-value <0.05 = coloured) with their effect direction (increased | decreased under aphid infestation (↑ | ↓) = brown | beige). A single metabolic feature (ID 9) was shaped by the interaction aphid infestation treatment × plant sex and is illustrated with one point for female (F) and male (M) plants. Numbered metabolic features correspond to Table 2. For further details see Supplementary Table S1A.

**Table 2:**
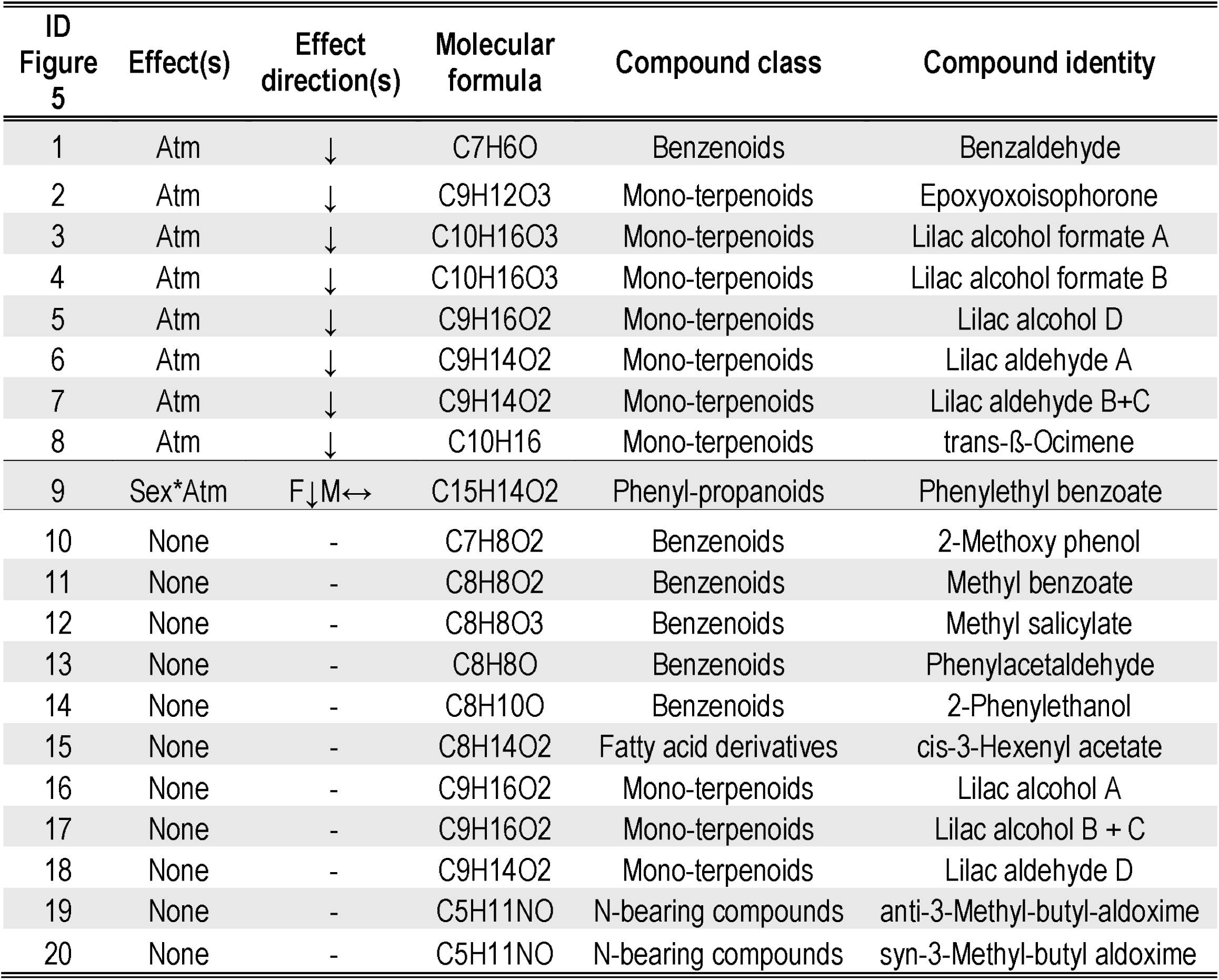
Overview of metabolites highlighted in Figure 5B. Metabolites highlighted with IDs in Figure 5 were significantly (uncorrected p-value < 0.05) and remarkably (log_2_ fold change <-1 or >1) modulated by aphid infestation treatment (atm; ↓ = reduced, ↑ = increased, - = unaffected by aphid infestation) and sex (F=female, M=male) and/or have been shown to elicit behavioural or antennal responses in the pollinator *Hadena bicruris* according to (Dötterl et al., 2006). Metabolites are ordered according to treatment effects, putative compound classes, and putative metabolite identities (alphabetically). See Supplementary Table S1A for a full list of floral VOC.

### 3.2 Effects of plant sex and aphid infestation on nectar traits

Nectar volume was unaffected by the aphid infestation treatment but considerably shaped by plant sex with female flowers featuring higher nectar amounts (*Table 1, Figure 6A, p< 0.001,* χ*²_1DF_= 46.10)*. The untargeted metabolome analyses of a standardized nectar volume per plant via UPLC-qTof-MS/MS yielded abundance data for 4,385 features. Of these, 761 features were tentatively identified via matches of their mass spectra with databases or via cheminformatics according (Hoffmann *et al*., 2022), and an additional 71 could be assigned to a compound class *(Supplementary Table 1B)*. This is the first detailed description of nectar chemistry in the ecological model system *S. latifolia.* The species’ nectar contained various nutritious metabolites, including sugars (D-glucose, D-fructose, sucrose, melezitose, stachyose, nystose, mannobiose, palatinose), proteinogenic and non-proteinogenic amino acids (e.g., proline, glutamic acid, tyrosine ornithine) and numerous oligopeptides, nucleosides (adenosine, adenosine monophosphate, a guanosine derivate), and multiple lipids (e.g., fatty alkyls; fatty acids, amides and alcohols; ceramides; glycerophosphoethanolamines, sphingolipids; prostaglandins). Its diverse secondary metabolites comprised vitamins (e.g., calcitriol, riboflavin), flavonoids (e.g., saponarin, vitexin-2’’-O-rhamnoside), terpenoids (e.g., myrtenol, perillyl alcohol, raddeanin D, pisumoside A), as well as fatty amides and amines (oleamide -a neuroactive compound [Mendelson and Basile, 2001] which has so far not been described in nectar, tyramine, homospermidine). Most of these secondary compounds are likely derived from plant metabolism, but several have a known fungal or bacterial origin and thus likely derive from the nectar microbiome (e.g., oleandomycin, enniatin B, asperiamide B, N-acetylglucosamine [chitin monomer]).

**Figure 6:**
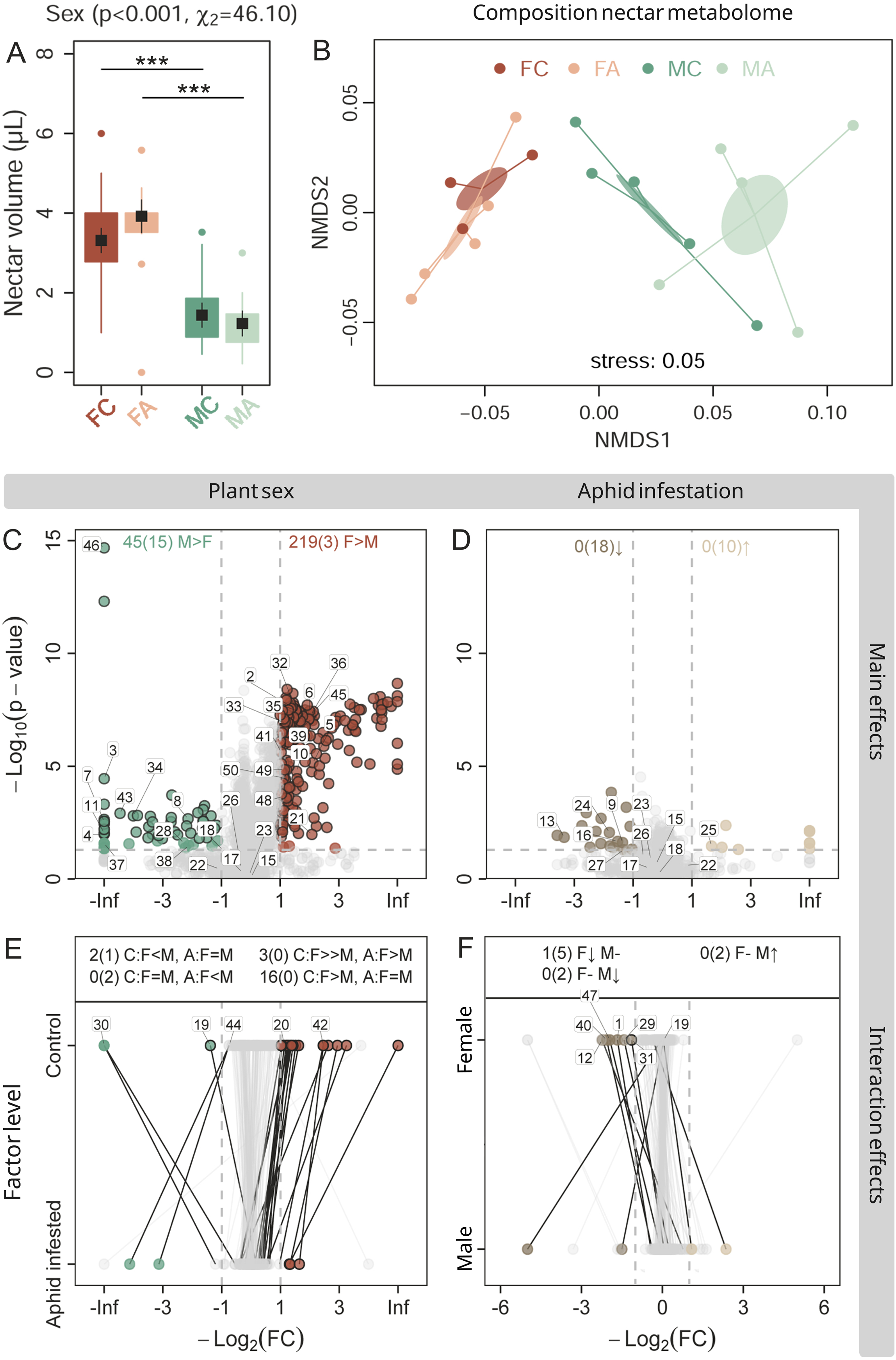
Combined effects of *Silene latifolia* plant sex and *Brachycaudus lychnidis* aphid infestation treatment on (A) nectar volume, and (B-F) the metabolic composition of a standardized nectar volume. (A) Box-Whisker plots of raw data for female control (FC, dark red, n = 23), female infested (FA, light red, n = 13), male control (MC, dark turquoise, n = 25), and male infested (MA, light turquoise, n = 25) plants with the marginal means +/- standard error (black squares) for these groups estimated by the (G)LMMs. Significant effects detected via type III ANOVAs based on Wald χ² tests are denoted at the top of the plot, while the results of relevant post-hoc comparisons are denoted within the plots (significance levels: ***p<0.001, **p<0.01, *p<0.05). (B) The NMDS ordination for FC (n = 5), FA (n = 3), MC (n = 5), and MA (n = 5) plants shows plant individual scores (points), distances to group centroids (spiders), and their standard error (ellipses).(C-F) Volcano plots illustrate the results of model loops, i.e., the main and interactive effects of plant sex and aphid infestation treatment on the abundance of individual metabolic features. Modulated features are highlighted (FDR corrected [uncorrected] p-value <0.05 and log_2_ fold change <-1 or >1 = coloured with black margin [coloured without margin]) with their effect direction (plant sex: F>M = red, M>F = turquoise, aphid infestation treatment: increased under aphid infestation (↑) = brown, decreased (↓) = beige). Interaction effect plots adopt the same modulation colour coding for (E) the sex effect within control and aphid infested plants and (F) the aphid infestation effect within females and males. Identical features within these group comparisons are connected by lines. Numbered metabolic features correspond to Table 3. A maximum of 15 identified features was highlighted for each modulation direction. For further details see Supplementary Table S1B.

**Table 3:**
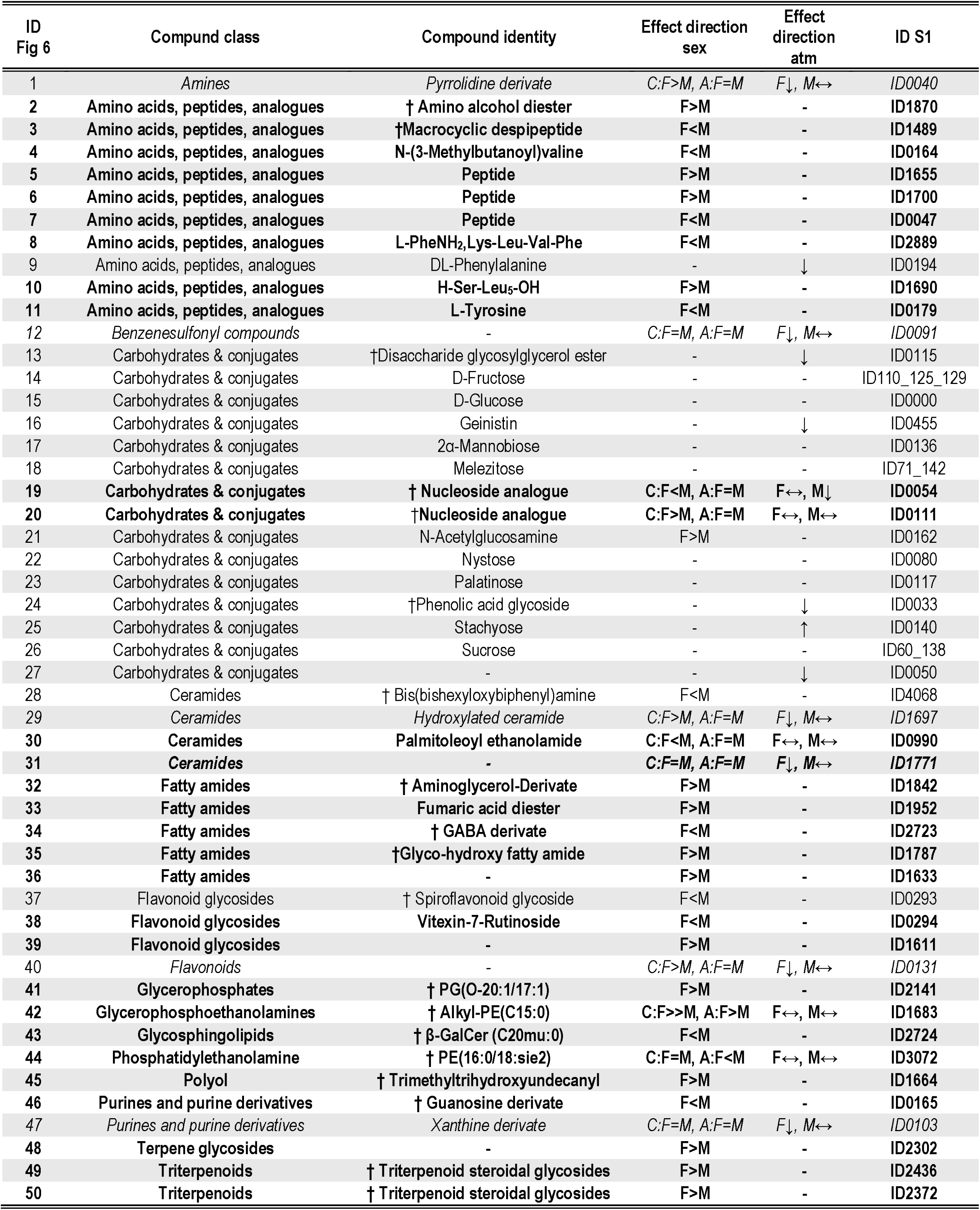
Overview of metabolites highlighted in Figure 6C-F. Metabolites highlighted with IDs in Figure 6C-F were saccharides (primary nutritional compounds) or putatively identified compounds that were significantly (FDR corrected [bold lettering] / uncorrected p-value <0.05) and remarkably (log_2_ fold change <-1 or >1) modulated by plant sex (F=female, M=male, entries only in column 2), aphid infestation treatment (atm; ↓ = reduced, ↑ = increased under aphid infestation, entries only in column 3) or the interaction plant sex × aphid infestation treatment (entries in column 2 and 3). The abundance of features printed in italic correlates with moth behaviour (Figure 3). Metabolites are listed with their compound class, putative identity, and ID in the full dataset Supplementary Table S1B. †: IUPAC names for the regarding metabolites are replaced with simplified names for compact presentation and fully listed S1B.

The PERMANOVA detected a highly significant effect of plant sex *(Table 1, p < 0.001, F_1DF_ = 10.65, R² = 0.39)*, but no effects of infestation treatment or the interaction sex × infestation treatment on the composition of the metabolome in a standardized volume of nectar. The group dispersions visualized in ordination plots via NMDS illustrate a clear differentiation of the nectar metabolome among female and male plants and additionally indicate different responses to the aphid infestation treatment in males and females in some features *(Figure 6B)*. The results from the linear model loop further support this observation *(Figure 6C-F, Table 2, Supplementary Table S1B).* When accounting (not accounting) for multiple testing, a total of 219 (3) features had significantly and remarkably (log_2_ fold change >1) higher abundance in females than males, including six features that were absent in the latter. These did not include primary nutritional compounds such as sugars or free amino acids, but seven oligopeptides, nine fatty amides, six terpenoids (including three triterpenoid steroids), a glycerophosphate, a flavonoid and the microbial compound N-acetylglucosamine [chitin monomer]. The remaining discriminatory features, including those unique to female nectar, could not be identified *(Figure 6C, Table 2, Supplementary Table 1B)*. On the other hand, 45 (15) features had significant and remarkably (log_2_ fold change <-1) higher abundances in males than females with 20 of them being absent in the latter. These included the free proteinogenic amino acid tyrosine (unique), N-(3-Methylbutanoyl)valine (unique), two oligopeptides (one unique), a macrocyclic despipeptide of probable microbial origin (unique), two flavonoid glycosides (one unique), a glycerosphingolipid, and a guanosine derivate (unique, highly significant). The aphid infestation treatment had clearly weaker effects on the composition of plant nectar. When accounting (not accounting) for multiple testing, plants showed increased abundances of 0 (10) features including the tetrasaccharide stachyose; and decreased abundances of 0 (18) features including the amino acid phenylalanine, a disaccharidester, and the isoflavonoid genistin *(Figure 6D, Table 2, Supplementary Table S1B)*. The interaction plant sex × aphid infestation treatment shaped the abundance of 21 (7) features *(Figure 6E& F; please note that both plots display the same metabolic features with different group comparisons; Table 2; Supplementary Table S1B)*. These interactions had seven major effect directions, which are listed as labels at the top of both plots. The most prominent interaction pattern was significantly and markedly higher metabolite abundance in females compared to males under control conditions, but not under aphid infestation in 16 (0) features, which included a nucleoside analogue. The second most prominent interaction pattern correlated with moth behaviour. In total 1(5) features had significantly and markedly reduced abundances under aphid infestation in females, but not in males. We analysed the MS² spectra for these six features in detail. Most of them did not yield high confidence matches or similarity scores to known compounds, suggesting these metabolites are not previously reported natural products. Our detailed analyses provided chemical class annotations and further details on structural features allowing to discuss their putative function in the context of plant-pollinator interactions (*Supplementary Table S1C*). These compounds include a pyrrolidine derivative (C₁₇H₁₈N, m/z 237.021) with aromatic substituents (m/z = 70.0648 ➔ pyrrolidinegroup, m/z = 77.0382 ➔ phenylmethylgroup, m/z = 91.0539 ➔ phenylgroup), a xanthine derivative with additional substituents (C₁₄H₁₄N₄O₅S, m/z 351.07), a benzenesulfonyl compound with a sugar-like substituent (C₁₄H₂₁NO₇S, m/z 351.07), two ceramides of which one bared multiple amino substituents (C₃₂H₆₅NO₄, m/z 528.3914; C₂₃H₅₁N₅O₄, m/z 484.3645) and a flavonoid (C₃₁H₂₄O₉, m/z 541.1492).

## 4. Discussion

Building on the pronounced sexual differentiation in floral traits of the dioecious plant *Silene latifolia*, we examined how controlled infestation by the oligophagous aphid *Brachycaudus lychnidis* affects interactions with the specialist moth pollinator *Hadena bicruris*. Contrary to classical expectations from sexual selection theory, female plants were not outcompeted by males in floral attractiveness under control conditions. Instead, they exhibited high plasticity in floral trait expression and produced diverse, potentially bioactive nectar metabolites, some of which may specifically target female pollinator behaviour. While aphid infestation affected some floral traits equally in both sexes and others more strongly in males or in females, we observed stronger declines in female attractiveness to *H. bicruris*. This points to sex-specific plastic responses, such as increased attraction of alternative pollinators in females, as well as sex-specific defence strategies, which we discuss here in detail.

### 4.1 Female *Silene latifolia* plants challenge expectations: no evidence of reduced floral attractiveness compared to males

Bateman’s theory of sexual selection predicts that, relative to males, female reproductive success is more limited by available resources and therefore does not increase with the number of mates, leading to less attractive flowers that serve as a resource exploited and competed for by males (Bateman, 1948). In the present study, we found no evidence for these predictions.

*Hadena bicruris* moths did not generally prefer single flowers of either sex (*Figure 4B*, *Table 1* - pollinator choice F-M). Female and male plants exhibited no general differences in flower numbers (*Figure 4A* - control conditions), flower sizes (*Table 1* - petal area), and flower colours (*Figure 4C*). Female flowers even featured higher nectar volumes as compared to males (*Figure 6A*) and major nectar sugar constituents showed no sex-specific differences (*Figure 6C*, *Table 3*, *Supplementary Table S1B*). Instead, we observed higher abundances of over 200 other primary and secondary metabolites in females than males, including six that were unique to female nectar. Some of these compounds likely shape the nutritional and pharmacological value of nectar for moth pollinators. These included seven oligopeptides, partially enriched in branched-chain amino acids, which may serve as a rapid source of NADH, FADH₂, and acetyl-CoA for ATP production, supporting energy demands during flight or allocation to egg production in female moths (Levin *et al*., 2017; Cavigliasso *et al*., 2023); nine fatty amides potentially serving as energy stores, metabolic fuel or behavioural modulators (Ranger *et al*., 2005; Yoshinaga, 2016; Levin *et al*., 2017); six terpenoids (of which two were steroidal glycosides) that may provide selective feeding or oviposition cues (González-Coloma *et al*., 2011; Boncan *et al*., 2020); a glycerophospholipid that may support the maintenance and generation of somatic tissue due to its central role in rapid membrane synthesis (Toprak *et al*., 2020); and a flavonoid which may act as health promotor in moth pollinators (Palmer=Young *et al*., 2019). The function and further identification of the numerous female biased metabolites, specifically the unidentified unique ones, will be the subject of future studies. As only female plants bear the cost of the interaction with *H. bicruris* (fruit predation), and only female moths impose this cost, there is great potential for the discovery of nectar metabolites interfering with or stimulating reproduction in female moths. Male *S. latifolia* plants, on the other hand, exhibited higher abundances of over 40 compounds, including several from the same chemical classes and with similar putative functions as those dominant in female nectar (BCAA-rich oligopeptides, flavonoid glycosides, membrane building lipids such as ceramides and glycosphingolipids). Most strikingly, tyrosine and a derivate of guanosine were unique to male nectar. Both compounds can be considered essential for the nutrition of moth pollinators that rely exclusively on nectar as a nitrogen source and may enhance its taste and memorability accordingly (Nepi, 2014). In addition, tyrosine is a precursor for the synthesis of the neurotransmitters tyramine and dopamine, which shape numerous insect behavioural traits like e.g., motivation and learning (Barberis *et al*., 2023). To the best of our knowledge, our study reports one of the most extensive cases of sexual differentiation in the composition of nectar for a dioecious plant to date (see also Nepi *et al*., 2001; Dötterl *et al*., 2014). Since *H. bicruris* showed no preference for either plant sex at the single-flower level (*Figure 4B*, *Table 1* - pollinator choice F-M), we conclude that these striking differences complement each other functionally to stabilize a dioecious system relying on frequent visits to both sexes, with plants promoting this by offering essential nectar resources in only one sex.

Overall, our study offers a more nuanced view of female floral traits than predicted by Bateman’s principle. A potential criticism is that females in our greenhouse setting did not exhibit lower floral attractiveness as they were not pollinated and hence did not invest in seeds, which aligns with long-term personal observations of visual flower traits in *S. latifolia*. However, this very aspect underscores the limitations of Bateman’s theory as it supports that females exhibit adaptive plasticity in floral traits, with attractiveness decreasing only as pollination success increases. High attractiveness of unpollinated female plants may promote female mate competition, post-pollination mate choice in females (Tonnabel *et al*., 2021), multiple paternity in fruits, and ultimately greater offspring diversity and fitness (Teixeira and Bernasconi, 2007). This would be in line with the theory of flexible sex role behaviour (Gowaty and Hubbell, 2005), which -similar to Bateman’s principle- originates from zoology and may be applicable to understanding reproductive strategies in dioecious plants. The substantial among-population variation in sexual dimorphism observed in *S. latifolia* in previous studies (Dötterl *et al*., 2009; Schrieber *et al*., 2017, 2019, 2021) further contradicts the assumption of a fixed evolutionary trajectory, instead pointing toward locally driven and context-dependent trait evolution.

### 4.2 Aphid infestation reduces female floral attractiveness via sex-specific trait changes

In accordance with our hypothesis, infestation with *B. lychnidis* aphids affected a wide range of traits in *S. latifolia* flowers, as well as the behavioural responses of *H. bicruris* moth pollinators. Although most of the observed effects clearly depended on plant sex, we begin with a general discussion of aphid-induced changes, as these have, to our knowledge, not been previously reported.

Aphid infestation reduced flower numbers (more strongly in females than males, *Figure 4A*), and reduced flower light reflection in the blue and green range (only in males, *Figure 4C*). These adverse effects likely result from both resource depletion and the reallocation of resources toward plant defence (Twayana *et al*., 2022), and may substantially limit plant visibility to pollinators (Barbot *et al*., 2022). Aphid infestation also moderately reduced the emission of five key volatile organic compounds (VOCs) known to elicit antennal and feeding responses in *H. bicruris* (Dötterl *et al*., 2006), at the level of individual flowers (*Figure 5B*, *Table 2*). Nectar volume was not lower in aphid-infested plants (*Figure 6A*), likely because they produced fewer flowers and could consequently invest more nectar in each one. While the major sugar constituents remained largely unaffected (*Figure 6D*, *Table 3*), stachyose levels increased in both sexes, suggesting that plants load their phloem with this poorly digestible tetrasaccharide to impair aphid performance (Twayana *et al*., 2022), while simultaneously ensuring pollination by enzymatically breaking it down into simpler sugars such as fructose, glucose, or sucrose in nectar. Conversely, levels of phenylalanine, a nucleoside analogue, a phenolic acid glycoside, and an isoflavonoid were reduced. These changes may result either directly from phloem-feeding-induced resource depletion or reflect a passive carry-over of defensive phloem sap modifications targeting aphids (Twayana *et al*., 2022). Although most of these alterations affected both sexes and would be expected to generally reduce pollinator attraction, this was observed only in one sex.

At the single-flower level, *H. bicruris* preferred flowers from control plants over aphid-infested ones only in *S. latifolia* females, but not males (*Figure 3*A). These preferences were also dependent on moth sex, being primarily exhibited by female *H. bicruris* (*Table 1*). This behaviour coincided with a reduction of the floral VOC phenylethyl benzoate - a compound not previously described in *S. latifolia* but found in other moth-pollinated species (El-Sayed, 2011) - which was observed only in aphid-infested female plants, but not in males (*Figure 5B*, *Table 2*). Additionally, six metabolic features in the nectar of *S. latifolia* were reduced by aphid infestation only in female plants (*Figure 6F*, *Table 3*). Four of them simultaneously had higher abundances in female than male plants only under control but not under aphid infestation conditions (*Supplementary table S1B*). Although these effects were only observed before FDR correction, the corresponding effect direction in pollinator behaviour suggests biological relevance. None of these metabolites belonged to the major nutritional nectar constituents (sugars, amino acids) or could be clearly identified as a previously described natural product. Our detailed chemoinformatic analysis of MS^2^ spectra elucidated compound classes and further structural components indicating a potential role of these metabolites as selective feeding cues or behavioural modulators for female *H. bicruris*. One candidate compound contains both pyrrolidine and phenyl substituents, a structural motif found in several synthetic psychoactive compounds that inhibit monoamine reuptake transporters, particularly dopamine, and are associated with stimulant activity (Meyer *et al*., 2013). Another notable candidate is a xanthine derivative, a class of compounds widely documented to enhance learning, memory, and arousal in pollinators (Mustard, 2020). The benzenesulfonyl compound containing amino and methoxy sugar-like moieties, the two ceramides and the flavonoid may possibly contribute to specific recognition by pollinators via contact cues. A detailed structural characterization of these novel natural products via NMR analysis of larger nectar volumes as well as individual and combinatorial bioassays are necessary to validate these putative ecological functions.

Even if more than just a small subset of the many measured floral traits had declined more severely upon aphid infestation in females than in males (as was in fact not the case), it would remain an overly simplistic conclusion that females are more vulnerable to stress. It remains to be clarified which of these responses represent a direct negative consequence of stress and which reflect a functional plastic adjustment under natural conditions. The finding that the nursery pollinator *H. bicruris* avoided flowers of aphid-infested *S. latifolia* females only *(Figure 3)*, combined with the greater choosiness of female moths *(Supplementary Figure S2)*, suggests that, under field conditions, infested female plants could experience a shift in the outcome of the interaction towards the latter end of the seed predation-pollination continuum. Additionally, reduced female flower attractiveness to *H. bicruris*, potentially mediated by shifts in nectar metabolite composition, could be associated with increased reliance on or even attraction of other pollinators that are less efficient but also less damaging. Moreover, the role of sex-specific defence strategies against aphids deserves particular attention in future studies. Our approach bypassed aphid host preferences; direct plant defences preventing the early establishment of aphids, the attraction of aphid parasitoids and predators that all occur in the field. Indeed, female plants in some dioecious species have been shown to be less frequently attacked by herbivores (Sargent and McKeough, 2022), to investigate more into direct defence (Sargent and McKeough, 2022) and potentially also into indirect defences (Aranda-Rickert *et al*., 2021). In fact it has been shown that *S. latifolia* females in Europe are clearly less frequently attacked by aphids than males (Wolfe *et al*., 2004). Hence, they are unlikely to experience aphid infestation as strong as in our experiment under more natural conditions. Yet, the cause for lower aphid abundances on female plants remains to be investigated in the future. Sex-specific indirect defences against aphids appear to be a particularly promising research route here. The seed-predating larvae of *H. bicruris* are attacked by a broad range of parasitoids and predators (Villacañas de Castro and Hoffmeister, 2020) that -if the plant is involved at all-should be attracted by females only. There is ample evidence that the evolution of indirect defences against different enemy species and guilds is tightly coupled via shared plant signalling and VOC synthesis pathways or the attraction of generalist predators (Kessler and Baldwin, 2001; Heil and McKey, 2003; Rasmann *et al*., 2005).

## 5. Conclusion and outlook

Our study provides evidence for multidirectional sexual differentiation of visual and chemical flower traits and their responses to aphid infestation in a dioecious plant species. It supports that the expression of sexual dimorphisms in floral attractiveness can be highly plastic depending on the presence of both, pollinators and herbivores, thereby opening exciting avenues for future research. Controlled experiments manipulating the presence of both, pollinators and aphids, ideally under field conditions reflecting natural conditions, while integrating data collection on plant, pollinator and aphid reproductive success with quantitative leaf and flower chemistry are a next step in disentangling the sex-specific co-evolution in this tripartite system. A particular focus will be on the many newly discovered sex-specific nectar metabolites that potentially serve as oviposition cues, physiologically reduce fecundity or offspring survival, or force visits to both plant sexes. In such future studies, it might be more purposeful to focus on the complementarity of female and male defence and floral attractiveness traits in stabilizing a dioecious reproductive system involving a nursery pollinator and several other parties (aphids, aphid and caterpillar parasitoids and predators), instead of seeking for the better defended or more attractive sex.

## Acknowledgements

Tobias Möckel from the FabLab in Kiel developed a customised camera attachment for light filter switches that was used for the assessment of flower colour composition. Reza Esmaeilnia Talemi supported the collection of plant nectar for metabolome analyses and Meike Pfeiler prepared nectar extracts. Stefan Dötterl advised on the establishment of our *Hadena bicruris* laboratory population, and Eric Folz was involved in the maintenance of the aphid and pollinator laboratory populations. Roman Adler designed vector graphs for the study design illustrations; and Verena Zajonc and Katharina Grosser provided valuable technical support.

## 6. Supplementary data

Supplementary Table S1: Summary of floral metabolites in *Silene latifolia*: functional annotation and responses to sex and aphid infestation

Supplementary Figure S1: Representative images from the experimental procedure Supplementary Figure S2: Effects of moth sex on choice behaviour

## Author contributions

Conceptualization: K.S.; Methodology: K.S., T.S., E.K., S.U., W.B., and T.D.; Investigation: K.Z., T.S., and E.K.; Validation: K.S., T.S., E.K., and W.B.; Formal Analysis: K.S.; Resources:

T.S., E.K., A.E., S.U., T.D., and W.B.; Data Curation: K.S.; Writing – Original Draft

Preparation: K.S.; Writing – Review & Editing: K.S., E.K., W.B., S.U., T.S, T.D., and H.B.; Writing – Manuscript Revision: K.S., and E.K.; Visualization: K.S.; Supervision: K.S., S.U.; Project Administration: K.S.; Funding Acquisition: K.S.

## 9. Data availability

All data supporting this article are be deposited in dryad under the following link: https://doi.org/10.5061/dryad.83bk3jb6q. The code used for statistical analyses and generation of graphs is deposited in zenedo under the following link: https://doi.org/10.5281/zenodo.17700170.

## Abbreviations used in the main text

ANOVA: analysis of variance
FC, FA, MC, MA: female control, female aphid infested, male control, male aphid infested
GC-MS: gas chromatography-mass spectrometry
(G)L(M)M: (generalized) linear (mixed effects) model
IQR: Interquartile range
NMDS: nonmetric multi-dimensional scaling
UHPLC-ESI-qTof-MS/MS: quadrupole time of flight - electro spray ionization - tandem mass spectrometry
PERMANOVA: permutational analysis of variance
Q1/3: Quartile 1/3
VOCs: Volatile organic compounds

