## Supplementary Figures S1 and S2 for "Aphid infestation induces plant-sex-specific changes in floral chemistry and pollinator behaviour in *Silene latifolia*"

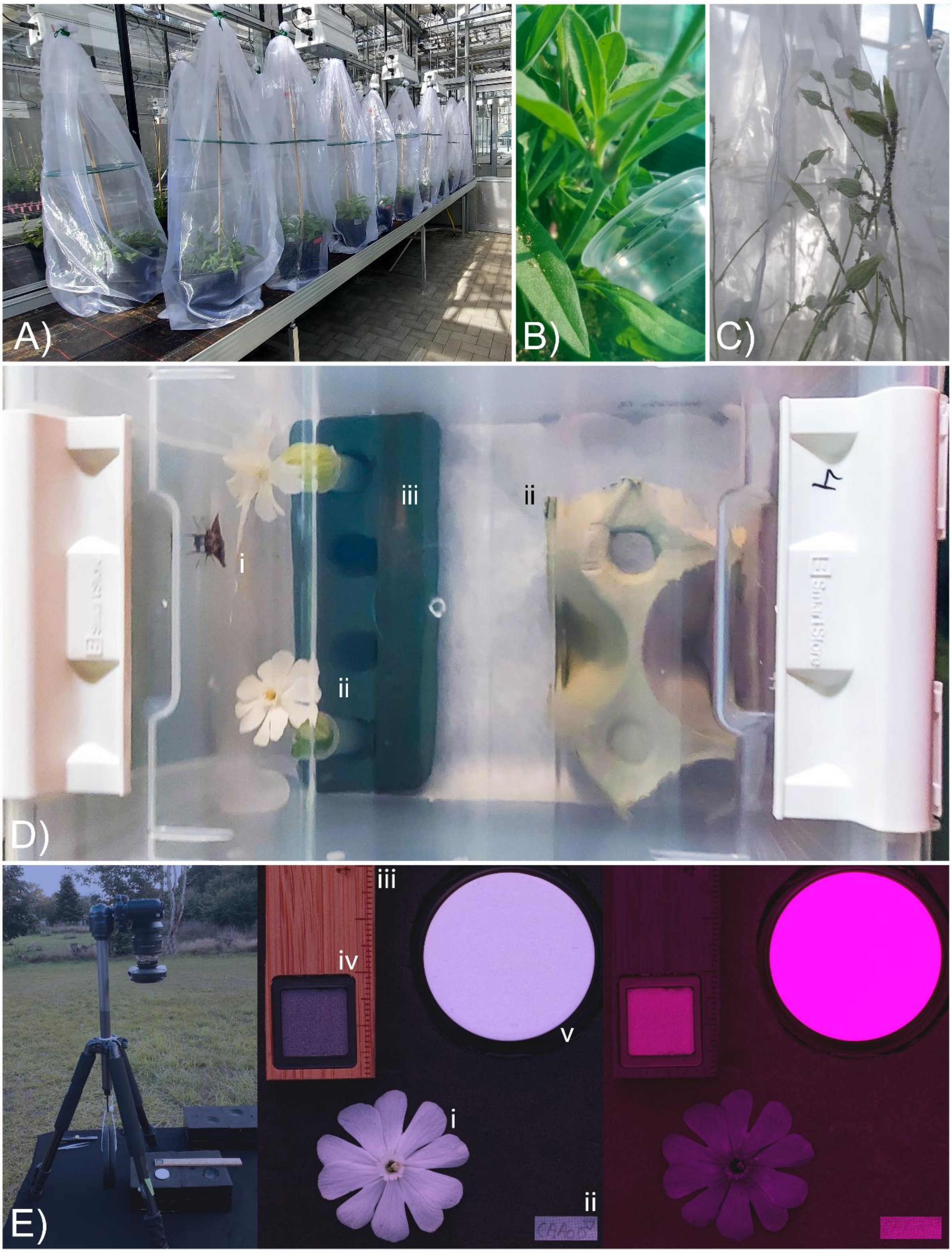


**Supplementary Figure S1:** Insight into the experiment. A - C) Experimental setup for the controlled aphid infestation. A) all plants were enclosed in 150 × 50 cm organza mesh bags, B) half of the bagged plants were then inoculated with 50 aphids each, and C) these aphids successfully established colonies. D) Pollinator choice assays. A single moth (i) was kept in a transparent and ventilated 15×12×11 cm PET box and was provided with a (ii) shelter and a (iii) feeding station equipped with two individual flowers, which were offered in water-filled Eppendorf tubes (v). E-F) Digital imaging setup. E) The camera was fixed on a tripod positioned on an exact horizontal platform, which was oriented towards the setting sun to take images of flowers in the F) visible light spectrum and G) the ultra-violet light spectrum. Images included an intact and fully opened flower (i) that was carefully plugged into a black ethylene vinyl acetate sheet equipped with a label (ii), a size standard (iii), a 10 % polytetrafluorethylene light standard (iv) and a 99 % spectralon light standard (v).

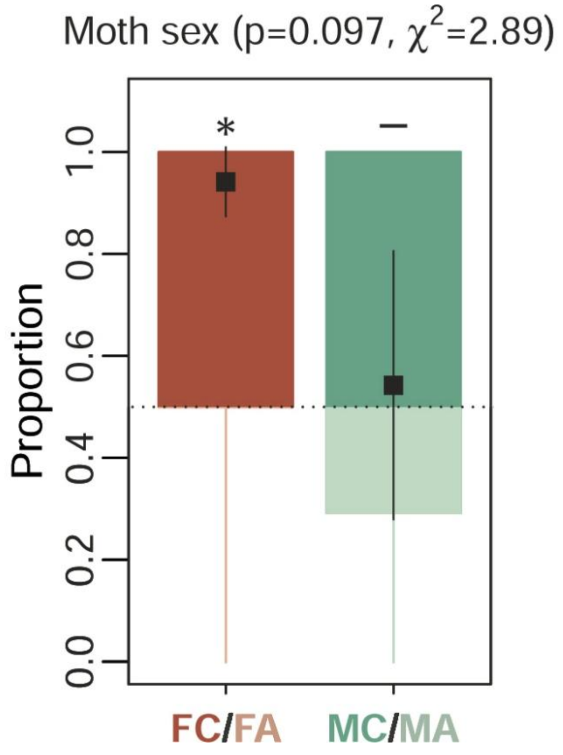


**Supplementary Figure S2:** Effects of *Hadena bicruris* sex on the choice behaviour analysed and illustrated as proportions of flower visits, feeding attempts and the total time spent on two offered *Silene latifolia* flowers (average over all behavioural parameters). The graph illustrates preference (i.e., deviation from a binomial outcome of 0.5 marked by the dashed line) for control plants (C, deep coloured, outcome > 0.5) versus aphid infested plants (A, light coloured, < outcome 0.5) as average over both plant sexes in female moths (F, red) and male moths (M, turquoise) (FC/FA: n=39, MC/MA: n=60). Graphs combine Box-Whisker plots of raw data (box: IQR Q1-Q3, whiskers: smallest/largest values within Q1/Q3 -/+ 1.5 × IQR, points: outliers) with the marginal means +/- standard error (black squares with black lines) estimated for all groups by the (G)LMMs. Effects assessed via type III ANOVAs based on Wald $\chi^2$ tests are denoted at the top of the plot, while the results of relevant post-hoc comparisons within plots indicate significant deviations of the binomial outcome from 0.5 (***p<0.001, **p<0.01, *p<0.05, - not significant).
